# Transcriptional Control of Brain Tumour Stem Cells by a Carbohydrate Binding Protein

**DOI:** 10.1101/2021.04.14.439704

**Authors:** Ahmad Sharanek, Audrey Burban, Aldo Hernandez-Corchado, Ariel Madrigal, Idris Fatakdawala, Hamed S Najafabadi, Vahab D Soleimani, Arezu Jahani-Asl

## Abstract

Brain tumour stem cells (BTSCs) and intratumoural heterogeneity represent major challenges in glioblastoma therapy. Here, we report that the *LGALS1* gene, encoding the carbohydrate binding protein, galectin1, is a key regulator of BTSCs and glioblastoma resistance to therapy. Genetic deletion of *LGALS1* alters BTSC gene expression profiles and results in downregulation of gene sets associated with mesenchymal subtype of glioblastoma. Using a combination of pharmacological and genetic approaches, we establish that inhibition of *LGALS1* signalling in BTSCs impairs self-renewal, suppresses tumourigenesis, prolongs lifespan, and improves glioblastoma response to ionizing radiation in preclinical animal models. Mechanistically, we show that *LGALS1* is a direct transcriptional target of STAT3 with its expression robustly regulated by the ligand OSM. Importantly, we establish that galectin1 forms a complex with the transcription factor HOXA5 to reprogram BTSC transcriptional landscape. Our data unravel an oncogenic signalling pathway by which galectin1/HOXA5 complex maintains BTSCs and promotes glioblastoma.

## Introduction

Glioblastoma is the most common and aggressive primary tumour in the adult brain. The current standard of care includes surgical excision of the tumour followed by ionizing radiation (IR) and chemotherapy (Chen et al., 2012b; Stupp et al., 2005). Despite these intense treatments, the median survival rate for glioblastoma patients remains 16-18 months (Chinot et al., 2014; Stupp et al., 2005). Genetic and phenotypic heterogeneity in glioblastoma represent major challenges in therapy.

The Cancer Atlas Genome (TCGA) project has revealed that the receptor tyrosine kinases (RTK) are altered in 88% of the glioblastoma patients (Cancer Genome Atlas Research, 2008). Epidermal growth factor receptor (EGFR) gene amplification occurs in approximately 40% of glioblastoma (Furnari et al., 2007). In addition, 63% to 75% of the glioblastoma that over-express EGFR are also found to have rearrangements of the EGFR gene, resulting in tumours expressing both wild-type EGFR (wtEGFR) and mutated EGFR (Ekstrand et al., 1991; Ekstrand et al., 1992; Malden et al., 1988). The epidermal growth factor receptor variant III (EGFRvIII) is the most common EGFR activating mutation (Wikstrand et al., 1995). EGFRvIII leads to the induction of multiple oncogenic pathways including the activation of the transcription factor (TF) STAT3 (de la Iglesia et al., 2008). Despite the importance of EGFRvIII and STAT3 as therapeutic targets, compounds designed to suppress this pathway have not yet led to promising outcomes in clinic (Raizer et al., 2010; Reardon et al., 2015). Therefore, identification of EGFRvIII/STAT3 dependent pathway(s) and drug targets in combination with complementary approaches is a promising avenue of research.

Glioblastomas are cytogenetically heterogeneous tumours that frequently display chromosomal copy number alterations including the gain of whole chromosome 7. Recently, the gene encoding the TF, Homeobox A5 (HOXA5), was reported to promote selection for the gain of chromosome 7 in human and mouse gliomas (Cimino et al., 2018). Importantly, the expression of HOXA5 correlated significantly with a more aggressive phenotype of glioblastoma (Cimino et al., 2018). The mechanisms that control HOXA5 function in glioblastoma remain largely unknown. Furthermore, the role of HOXA5 in EGFRvIII subtype of tumour is uncharacterized.

The lactose binding lectin, galectin1, encoded by the *LGALS1* gene, is a member of the carbohydrate binding proteins defined by their ability to recognize beta-galactose molecules found on cell surfaces and extracellular matrices (Liu and Rabinovich, 2005). Recent studies have found that galectin1 is highly expressed in different human cancers including colon, breast, lung, head and neck, ovarian, prostate, and gliomas (Astorgues-Xerri et al., 2014a). The upregulated expression of *LGALS1* in high-grade glioma correlates significantly with poor patient prognosis (Rorive et al., 2001). Higher extracellular galectin1 levels have been also detected in the plasma of glioblastoma patients (Kros et al., 2015). Several findings highlight that galectin1 regulates glioblastoma tumour microenvironment and promotes migration (Camby et al., 2006), invasion (Toussaint et al., 2012), angiogenesis (Le Mercier et al., 2008), and immune escape (Verschuere et al., 2014). However, the role of *LGALS1* in cancer stem cells and in the genetic background of EGFRvIII, has remained unexplored.

Brain tumour stem cells (BTSCs) contribute to glioblastoma tumour heterogeneity by transitioning between different cell cycle states to: a) self-renew and sustain themselves, b) undergo persistent proliferation to contribute to tumour growth, c) differentiate and give rise to diverse cell populations to recapitulate the functional heterogeneity of the tumour, d) exit cell cycle and evade therapy, and e) re-enter cell cycle, and give rise to tumour recurrence (Bao et al., 2006; Chen et al., 2012a; Chen et al., 2012b; Galli et al., 2004; Lathia et al., 2015; Singh et al., 2004; Venugopal et al., 2015). Although these changes can stem from genetic alterations or response to environmental cues, the mechanisms that dictate BTSC fate are not fully understood. Defining the molecular mechanisms that govern BTSC fate in different glioblastoma subtypes is, therefore, critical for developing better treatments.

In the present study, we report that *LGALS1*/HOXA5 signalling pathway promotes BTSC self-renewal and glioblastoma tumourigenesis downstream of EGFRvIII/STAT3 signalling pathway. Importantly, *LGALS1* promotes mesenchymal subtype of glioblastoma and confers resistance of glioblastoma tumours to therapy.

## Results

### *LGALS1* is a direct transcriptional target of STAT3

EGFRvIII/STAT3 signaling is important in glioblastoma. To gain insights into EGFRvIII-gene signatures in patient-derived BTSCs, we analyzed public database in which RNA-seq analysis of human BTSCs that naturally harbour EGFRvIII mutation or lack the mutation, were employed to establish EGFRvIII candidate target genes (Jahani-Asl et al., 2016). *LGALS1* scored among the top candidate target genes with its mRNA significantly upregulated in multiple EGFRvIII-expressing BTSC lines, raising the question of whether *LGALS1* plays a role in the regulation of BTSCs that harbour EGFRvIII mutation. To begin with, we subjected patient-derived BTSCs to immunoblotting analysis and confirmed that the expression levels of galectin1 protein were significantly upregulated in human BTSCs harbouring the EGFRvIII mutation compared to control BTSCs lacking the mutation (**Figure 1a-b**). Next, to establish whether *LGALS1* is a downstream target of EGFRvIII, we induced knockdown (KD) of *EGFRvIII/EGFR* using a pool of siRNAs in EGFRvIII-expressing BTSCs and subjected the cell lysates to RT-qPCR (**Figure S1a**) and immunoblotting analyses (**Figure 1c-d**). Our data showed significant downregulation of *LGALS1* mRNA and galectin1 protein in *EGFRvIII* KD BTSCs (**Figure 1c-d and Figure S1a**). In parallel, we treated EGFRvIII-expressing BTSCs with lapatinib, an inhibitor of EGFR/EGFRvIII phosphorylation. Consistent with the results obtained with the genetic KD of *EGFRvIII*, pharmacological inhibition of EGFRvIII by lapatinib significantly reduced galectin1 protein expression level as revealed by immunoblotting and immunostaining (**Figure 1e-h and Figure S1b-c**). Interestingly, addition of the ligand EGF to BTSCs that do not harbour EGFRvIII induced the phosphorylation of wtEGFR and a robust increase in galectin1 protein level (**Figure S1d-f**). Our results established that EGFRvIII and phosphorylated EGFR induce the upregulation of *LGALS1* in patient-derived BTSCs.

**Figure 1.**
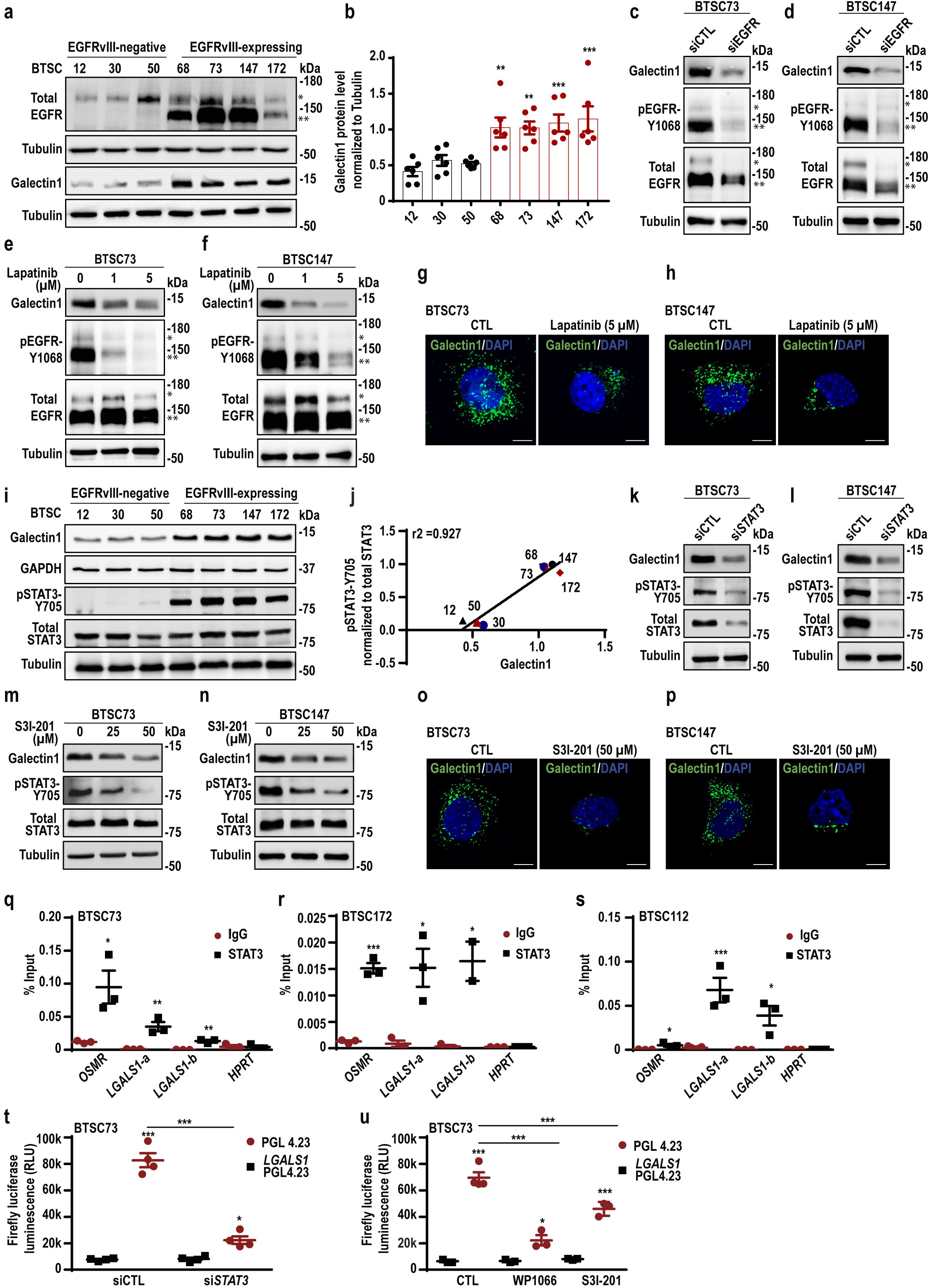
Galectin1 is a direct transcriptional target of STAT3 in patient-derived BTSCs. (**a**) BTSCs were subjected to immunoblotting analysis using the antibodies indicated on the blots. wtEGFR and EGFRvIII bands are marked with * and **, respectively. (**b**) Densitometric quantification of galectin1 protein level normalized to tubulin in different BTSC lines is shown. (**c-d**) *EGFR*/*EGFRvIII* KD (si*EGFR*) and control BTSCs (siCTL) were analyzed by immunoblotting as described in a. (**e-h**) BTSCs were treated with 1 or 5 µM lapatinib and galectin1 expression was assessed by immunoblotting (e-f) and immunostaining (g-h). Nuclei were stained with DAPI. Scale bar = 10 μm. (**i**) BTSCs were subjected to immunoblotting analysis using the antibodies indicated on the blots. (**j**) Pearson correlation analysis of pSTAT3-Y705 and galectin1 protein expression in different BTSCs is shown. (**k-l**) *STAT3* KD (si*STAT3*) and siCTL BTSCs were analyzed by immunoblotting as described above. (**m-p**) BTSCs were subjected to immunoblotting or immunostaining following treatment with 25 or 50 µM of the STAT3 inhibitor, S3I-201. Scale bar = 10 μm. (**q-s**) EGFRvIII-expressing BTSCs were subjected to ChIP using an antibody to STAT3 or IgG control followed by qPCR using two different pairs of primers (*LGALS1*-a and *LGALS1*-b). *OSMR*, and *HPRT* loci were used as positive and negative controls, respectively. (**t-u**) Luciferase reporter assay was performed in BTSC73 following KD of *STAT3* using siRNA (t) or treatment with STAT3 inhibitors, 5 µM WP1066 or 50 μM S3I-201 (u). Data are presented as the mean□±□SEM, n ≥ 3. Unpaired two-tailed *t*-test (q, r and s); one-way ANOVA followed by Dunnett’s test (b) or Tukey’s test (t and u),*p < 0.05, **p < 0.01, ***p < 0.001. See also Figures S1 and S2.

EGFRvIII and activated EGFR form a complex with the TF STAT3 to activate transcriptional networks that drive tumourigenesis in different human cancers including glioblastoma (de la Iglesia et al., 2008; Lo et al., 2005; Wheeler et al., 2010). For example, STAT3 directly occupies the promoters of *OSMR* and *iNOS* in EGFRvIII-expressing BTSCs and astrocytes (Jahani-Asl et al., 2016; Puram et al., 2012). Analysis of the human *LGALS1* promoter revealed the presence of multiple STAT3 consensus motifs (Schaefer et al., 1995; Seidel et al., 1995) (**Figure S2a**). Strikingly, in analysis of patient-derived BTSCs, we observed a positive correlation between the expression of galectin1 and phosphorylated STAT3 (Y705) (**Figure 1i-j**). We, thus, asked if EGFRvIII-mediated upregulation of *LGALS1* is controlled by STAT3. *LGALS1* expression levels were analyzed following pharmacological inhibition of STAT3 by WP1066 and S3I-201 or genetic KD of *STAT3* in EGFRvIII-expressing BTSCs (**Figure 1k-p and Figure S2b-c**). We observed a significant decrease in *LGALS1* mRNA and galectin1 protein level in *STAT3* KD BTSCs and BTSCs treated with either WP1066 or S3I-201 (**Figure 1k-p and Figure S2b-c**).

To determine whether endogenous STAT3 directly occupies the promoter of the *LGALS1* gene in EGFRvIII-expressing BTSCs, we performed ChIP-PCR and luciferase reporter assays in EGFRvIII-expressing BTSCs. In ChIP-qPCR analysis, we subjected three different human BTSC lines (#73, 112, and 172), to immunoprecipitation (IP) using an endogenous STAT3 antibody or an IgG control. Our analysis revealed a significant enrichment of endogenous STAT3 at the *LGALS1* promoter across all the BTSCs (**Figure 1q-s**). In analysis of the activity of the *LGALS1* promoter in transient expression assays, we found that the expression of a luciferase reporter gene that is controlled by 376 nt of the 5′ regulatory sequences of the *LGALS1* gene, was significantly downregulated upon deletion of *Stat3* in mouse astrocytes (**Figure S2d**). Consistent with these results, we found that KD of *STAT3* gene by siRNA or pharmacological inhibition of STAT3 by WP1066 and S3I-201 in BTSC73 significantly attenuated the expression of the *LGALS1*-luciferase reporter gene (**Figure 1t-u**).

Next, we examined if activation of STAT3 in BTSCs that lack the EGFRvIII mutation induces the expression of galectin1. We employed the ligand OSM, which robustly induces the phosphorylation of STAT3 via wtEGFR/OSMR complex in BTSCs (Jahani-Asl et al., 2016). OSM-mediated phosphorylation of STAT3 resulted in a robust increase in galectin1 expression level and KD of *STAT3* attenuated this effect (**Figure S2e-f**). Furthermore, OSM addition induced the binding of STAT3 to the promoter of *LGALS1* (**Figure S2g**). Taken together, we established that STAT3 directly occupies the promoter of *LGALS1* gene to upregulate its expression in EGFRvIII-expressing BTSCs. In BTSCs lacking the EGFRvIII mutation, however, addition of ligand is required to induce the phosphorylation of STAT3 and its binding to the *LGALS1* promoter.

### *LGALS1* promotes the growth and invasion BTSCs

EGFRvIII/STAT3 signalling plays crucial roles in cell growth and proliferation (Jensen et al., 2020; Stechishin et al., 2013). Identification of *LGALS1* as a direct downstream target gene of EGFRvIII/STAT3 oncogenic pathway in human BTSCs led us next to examine the role of *LGALS1* in the regulation of BTSC growth. We employed genetic approaches in which we induced KO or KD of *LGALS1* in EGFRvIII-expressing BTSCs using CRISPR-Cas9 or lentiviral-mediated transduction of BTSCs with shRNAs targeted against *LGALS1* mRNA, respectively (**Figure S3a-e**). We assessed cell proliferation by CellTiter-Glo luminescent assay in *LGALS1* CRISPR EGFRvIII-expressing BTSC lines (#73 and 147) and found a marked reduction in cell proliferation in *LGALS1* CRISPR BTSCs (**Figure 2a-b**). Consistent with the results of cell proliferation assay, cell population growth assay showed that deletion of *LGALS1* significantly suppressed BTSC growth (**Figure 2c**). Similarly, KD of *LGALS1* using lentiviral transduction with two different shRNAs targeting *LGALS1*, resulted in a significant reduction of BTSC population growth (**Figure S3f**). Interestingly, the decrease of cell proliferation upon deletion of *LGALS1* was not associated with an induction of cell death as analyzed by flow cytometry for annexin V and propidium iodide (PI) (**Figure S3g-h**).

**Figure 2.**
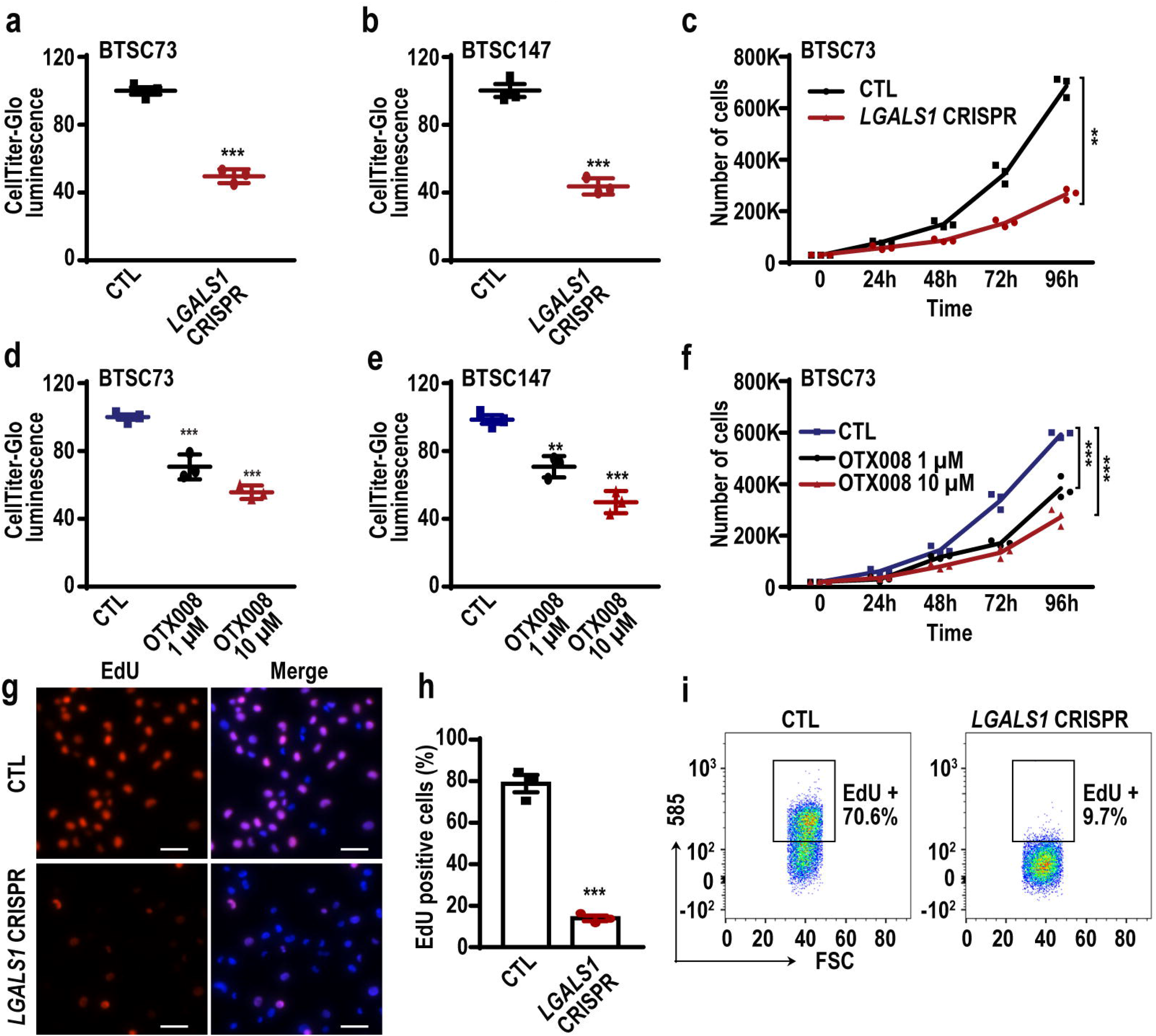
Galectin1 controls BTSC proliferation. (**a-b**) Cell viability was assessed by CellTiter-Glo assay in *LGALS1* CRISPR and CTL BTSCs. (**c**) Population growth curves for *LGALS1* CRISPR and CTL BTSC73 are shown. (**d-f**) Cell viability assay (d-e) and population growth curves (f) of BTSC73 treated with 1 or 10 µM OTX008 are shown. (**g**) Representative images of EdU staining in *LGALS1* CRISPR and CTL BTSC73 are shown. (**h**) The number of EdU positive cells was quantified using Fiji software. (**i**) EdU incorporation was analyzed by flow cytometry in *LGALS1* CRISPR and CTL BTSC73. Representative scatter plots of flow cytometry analyses are shown. Data are presented as the mean□±□SEM, n = 3. Unpaired two-tailed *t*-test (a, b, c and h); one-way ANOVA followed by Dunnett’s test (d, e and f), **p < 0.01, ***p < 0.001. See also Figures S3 and S4.

Next, we assessed the impact of OTX008, the pharmacological inhibitor of galectin1. OTX008 is a small molecule calixarene derivative which is designed to bind specifically to the galectin1 amphipathic β-sheets leading to its oxidation and proteasomal degradation (Astorgues-Xerri et al., 2014b). Treatment of BTSC73 and BTSC147 with OTX008 resulted in a dose-dependent inhibition of BTSC proliferation (**Figure 2d-f**). We confirmed that these differences were not due to toxicity or induction of apoptosis since 1-10 µM of OTX008 did not induce cell death as analyzed by annexin V/PI flow cytometry assay (**Figure S3i**). Furthermore, to confirm the role of *LGALS1* in BTSC proliferation, we performed 5-ethynyl-2’-deoxyuridine (EdU) assay. We observed a drastic reduction in the number of EdU-incorporating BTSCs in *LGALS1* CRISPR (**Figure 2g-i**) and OTX008-treated BTSC73 (**Figure S3j**).

Given that galectin1 is reported to be expressed at the tumor margin and regulates glioblastoma cell invasion (Toussaint et al., 2012), we next examined if galectin1 regulates the invasion of BTSCs. *LGALS1* CRISPR and CTL BTSC73 and 147 were subjected to collagen type I invasion assay and the invasion index was calculated. We found that silencing of *LGALS1* inhibits BTSC invasion in different EGFRvIII-expressing BTSC lines (**Figure S4a-d**). Altogether, our results demonstrate that *LGALS1* promotes growth, proliferation and invasion of BTSCs that harbour EGFRvIII mutation.

### Genetic knockdown of *LGALS1* or pharmacological inhibition of galectin1 impairs tumourigenic capacity of BTSCs

EGFRvIII/STAT3 signalling plays a key role in promoting glioblastoma tumourigenesis (de la Iglesia et al., 2008; Fan et al., 2013). Our *in vitro* data in which *LGALS1* impaired the growth and proliferation of BTSCs together with our discovery that *LGALS1* is a direct target of EGFRvIII/STAT3 signalling, raised the question of whether *LGALS1* regulates the growth of BTSC-derived tumours.

To begin with, we subcutaneously injected *LGALS1* CRISPR or control EGFRvIII-expressing BTSC73 into the flank of 8-week-old SCID mice and assessed the ability of these cells to form tumours *in vivo*. Three weeks following injection, all mice in the control group harboured ulcerated tumours at the site of injection and had lost more than 20% of their body weight at end point. Strikingly, no tumours were formed in mice group receiving the *LGALS1* CRISPR BTSCs (**Figure 3a-b**). Our data established that deletion of *LGALS1* impairs BTSC growth *in vivo*. Next, we assessed the impact of the pharmacological inhibition of galectin1 by OTX008 on tumour growth. We found that treatment of patient-derived xenografts with OTX008 significantly suppressed tumourigenesis in subcutaneous tumour assays performed with two different patient-derived BTSC lines (#73 and 147) (**Figure 3c-f**).

**Figure 3.**
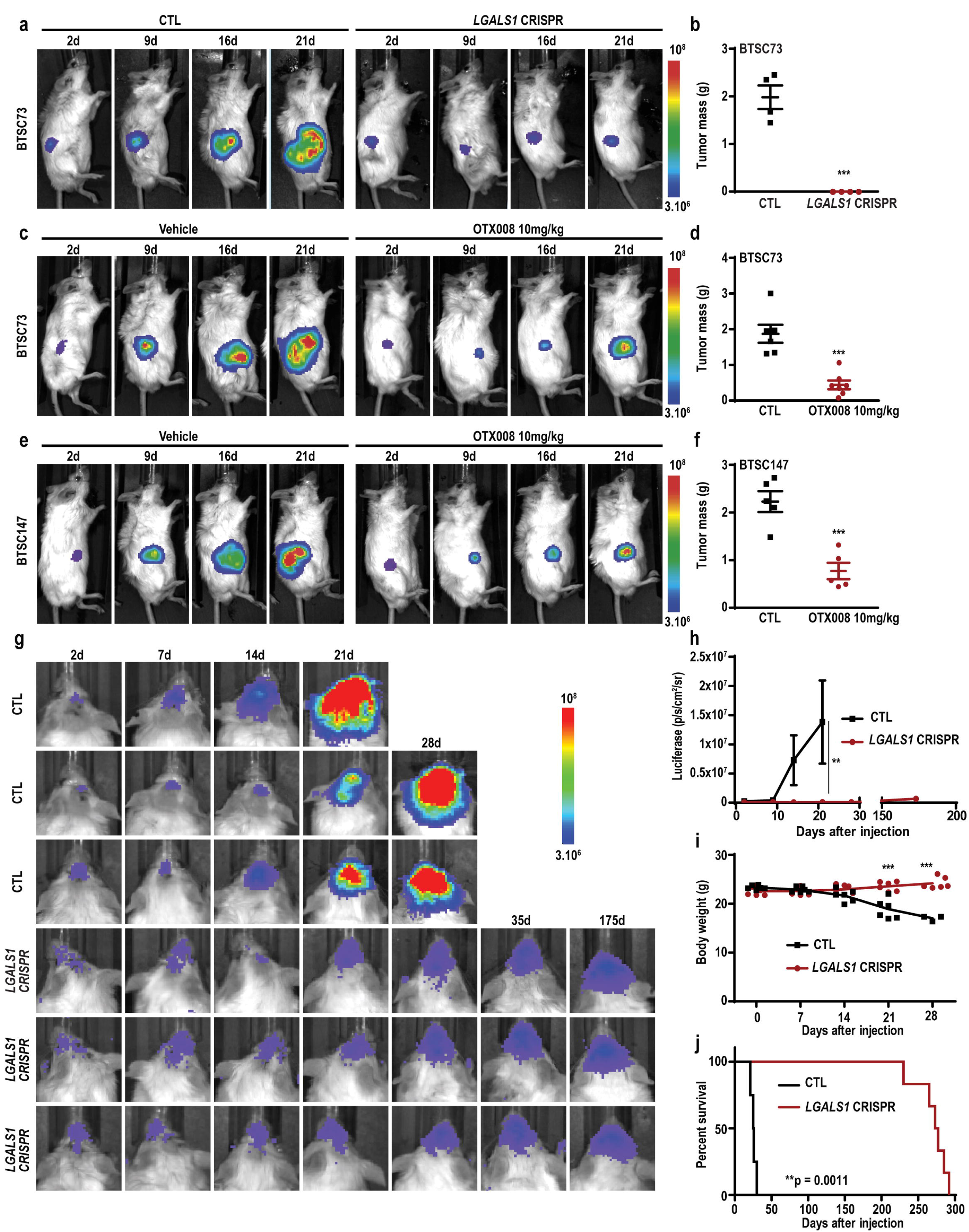
Galectin1 promotes glioblastoma tumourigenesis. (**a-b**) *LGALS1* CRISPR or CTL BTSC73 were subcutaneously injected into SCID mice. Representative bioluminescence real-time images tracing tumour growth are shown (a). Graph represents tumour mass (b). (**c-f**) BTSC73 or BTSC147 were injected subcutaneously into SCID mice and treated with 10 mg/kg OTX008. Representative bioluminescence real-time images tracing tumour growth are shown (c, e). Graphs represent tumour mass (d, f). (**g-j**) *LGALS1* CRISPR or CTL BTSC73 were intracranially injected into SCID mice. Representative bioluminescence real-time images tracing tumour growth are shown (g). Intensities of luciferase signal were quantified at different time points using Xenogen IVIS software (h). Graph represents quantification of animal weight (i). KM survival plot was graphed to evaluate mice lifespan in each group (j). Data are presented as the mean□±μSEM, n ≥ 4 mice. Unpaired two-tailed *t*-test (b, d, f, h and i); log-rank test (j), **p < 0.01, ***p < 0.001.

To investigate the impact of *LGALS1* deletion on brain tumour formation, we performed intracranial xenografts of *LGALS1* CRISPR or control EGFRvIII-expressing BTSC73 in SCID mice. Tumour volume was monitored via luciferase-based IVIS imaging (**Figure 3g-j**). At 21 days following surgery, mice receiving control BTSC73 formed brain tumours and were at endpoint as assessed by major weight loss and neurological signs (**Figure 3i**). Strikingly, no tumours were found in any of the animals receiving *LGALS1* CRISPR BTSC73. To investigate the impact of *LGALS1* deletion on the animal lifespan, we maintained the animals for up to one year and assessed survival. Kaplan-Meier (KM) survival plots revealed that deletion of *LGALS1* in BTSC73 expanded the lifespan of the animals by ∼9 months compared to only 21 days of mice bearing control BTSCs (**Figure 3j**). Our data established that *LGALS1* is a key regulator of tumourigenesis in brain tumours harbouring EGFRvIII mutation.

### Establishing *LGALS1* gene signatures in human BTSCs

Upstream of *LGALS1*, EGFRvIII/STAT3 signalling pathway functions to control the expression of *LGALS1* (**Figure 1, Figure S1-2**). To dissociate downstream signalling pathways controlled by *LGALS1*, we subjected the *LGALS1* CRISPR and control EGFRvIII-expressing BTSC73, to mRNA-seq analyses (**Figure S5**). Gene-expression profiling revealed differentially expressed genes in *LGALS1* CRISPR BTSCs compared to control (adjusted p-value < 0.1) (**Figure 4a**). We next assessed global gene expression changes in *LGALS1* CRISPR relative to control BTSCs by Gene Set Enrichment Analysis (GSEA). First, we sought to determine if silencing of *LGALS1* alters the gene signatures that define the TCGA transcriptional subtypes of glioblastoma (Verhaak et al., 2010). We found a significant downregulation of the mesenchymal subtype gene sets in *LGALS1* CRISPR (**Figure 4b**). This significant downregulation of mesenchymal gene set is associated with a tendency to the upregulation of the proneural gene signature (**Figure 4c**). Consistent with these results, analysis of the cell-state-specific signature genes (Neftel et al., 2019), revealed that loss of *LGALS1* led to downregulation of the hypoxia-independent mesenchymal-like meta-module (MES1-like) gene set (**Figure 4d**), a cell state that is enriched in the TCGA-mesenchymal subtype (Neftel et al., 2019). Furthermore, GSEA enrichment analysis showed a significant downregulation of candidate target genes that are involved in the regulation of the cell cycle, including G2/M transition and mitotic spindle assembly (**Figure 4e-f**). We validated the changes in the mRNA expression of select cell cycle candidate target genes in two *LGALS1* CRISPR BTSCs (#73 and 147) by RT-qPCR analysis (**Figure 4g-h**). Next, we examined the relevance of these transcriptional changes on the cell cycle regulation via performing cell cycle profiling by FACS analysis. Our data suggest that loss of *LGALS1* leads to an inhibition of cell cycle progression characterized by an accumulation of cells in the G2/M phase in multiple patient-derived EGFRvIII-expressing BTSCs (**Figure 4i-j**), highlighting a role for *LGALS1* as a positive regulator of the cell cycle in BTSCs.

**Figure 4.**
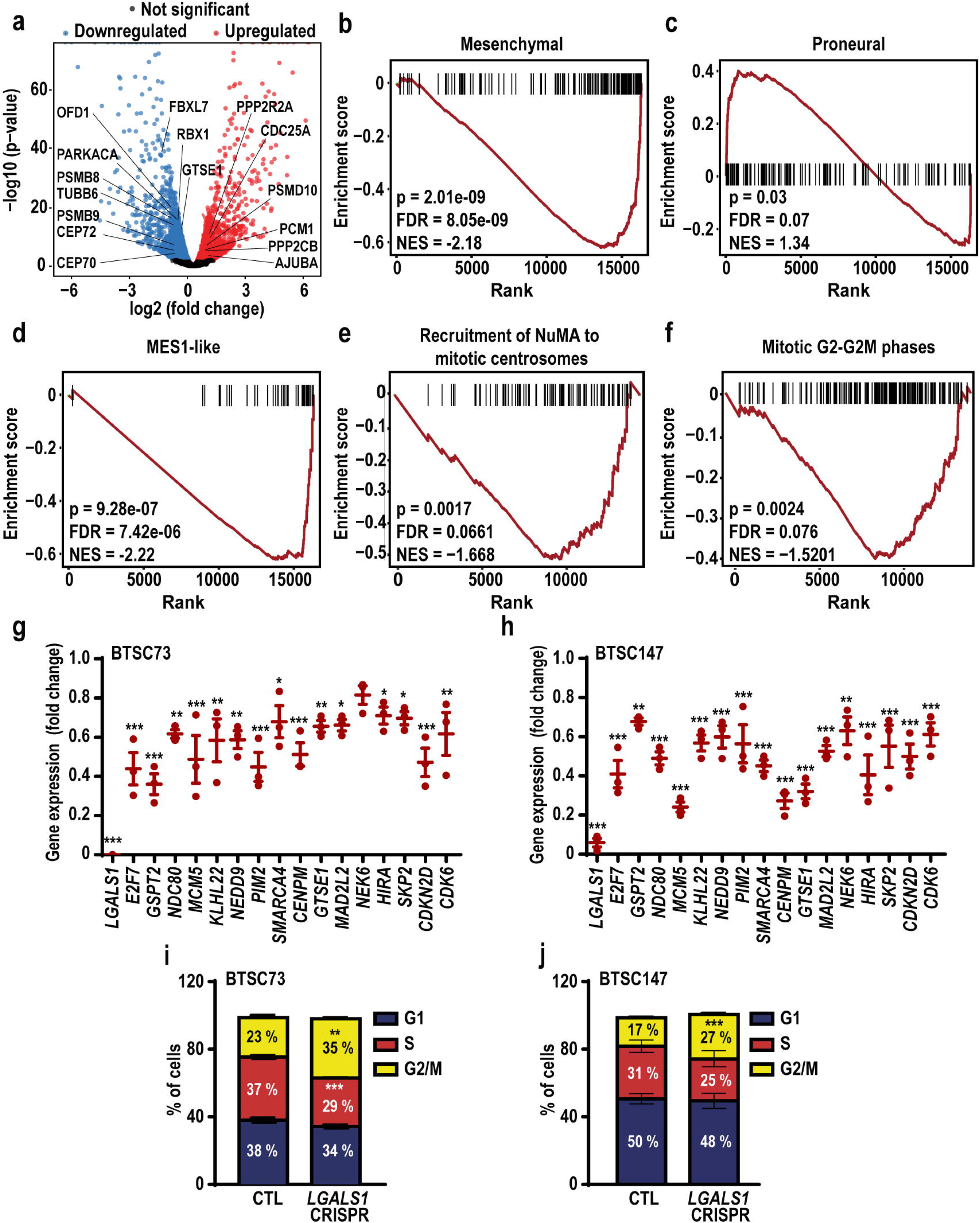
Genome wide analysis of *LGALS1*-differentially regulated genes. (**a**) Volcano plot representing *LGALS1* differentially regulated genes is shown. (**b-c**) GSEA analysis demonstrates enrichment for gene sets corresponding to mesenchymal (b) and proneural (c) subtypes of glioblastoma. (**d**) GSEA analysis demonstrates enrichment for gene sets corresponding to mesenchymal-like meta-module (MES1-like) signature. (**e-f**) GSEA analysis demonstrates enrichment for gene sets corresponding to recruitment of NuMA to mitotic centrosomes (e) and mitotic G2−G2/M phases (f). (**g-h**) RNA-seq data was validated by RT-qPCR in BTSC73 and BTSC147. (**i-j**) Cell cycle distribution was assessed by flow cytometry after PI staining in *LGALS1* CRISPR BTSCs. Data are presented as the mean□±□SEM, n = 3. One-way ANOVA followed by Dunnett’s test (g and h); unpaired two-tailed *t*-test (i and j), *p < 0.05, **p < 0.01, ***p < 0.001. See also Figure S5.

### *LGALS1* regulates BTSC self-renewal

Cancer stem cells hijack transcriptional and epigenetic programs that endow them with a unique feature of long-term self-renewal potential, resulting in the continuous expansion of self-renewing cancer cells and tumour formation (Brooks et al., 2015; Kreso and Dick, 2014). Cell cycle control is crucial for maintenance of stem cells, and disruption of cell cycle regulators has been shown to impair stem cell self-renewal (He et al., 2009), raising the question of whether *LGALS1* regulates the self-renewal capacity of BTSC. To examine the impact of *LGALS1* on self-renewal, we subjected *LGALS1* CRISPR and control EGFRvIII-expressing BTSCs to limiting dilution assay (LDA) and extreme limiting dilution assay (ELDA) (Hu and Smyth, 2009). Strikingly, deletion of *LGALS1* resulted in a significant decrease in sphere numbers and sphere-formation frequency compared to corresponding controls (**Figure 5a-d**). In addition, clonogenic assay was performed by seeding 1 cell per well and we found that loss of *LGALS1* resulted in a significant decrease in sphere forming capacity (SFC) of different EGFRvIII-expressing BTSCs (**Figure 5e-f**). Consistent with this data, KD of *LGALS1* by lentiviral transduction of two different shRNAs (**Figure S6a-b**) or transient KD of *LGALS1* using *a* pool of siRNAs (**Figure S6c-d**), significantly decreased BTSC self-renewal. Given that *LGALS1* is upregulated in an EGFRvIII/STAT3-dependent manner, we sought to examine whether *LGALS1*-mediated regulation of BTSC self-renewal is specific to EGFRvIII cohort of BTSCs. We, therefore, induced the KD of *LGALS1* in two patient-derived BTSCs that do not harbour EGFRvIII mutation (#12 and 30) and subjected the cells to LDA and ELDA analyses. Our results demonstrated that KD of *LGALS1* had no significant impact on the number of spheres (**Figure S6e-f**) or the frequency of sphere-formation (**Figure 5g-h**), perhaps due to low expression levels of *STAT3/LGALS1* in BTSCs lacking the EGFRvIII mutation.

**Figure 5.**
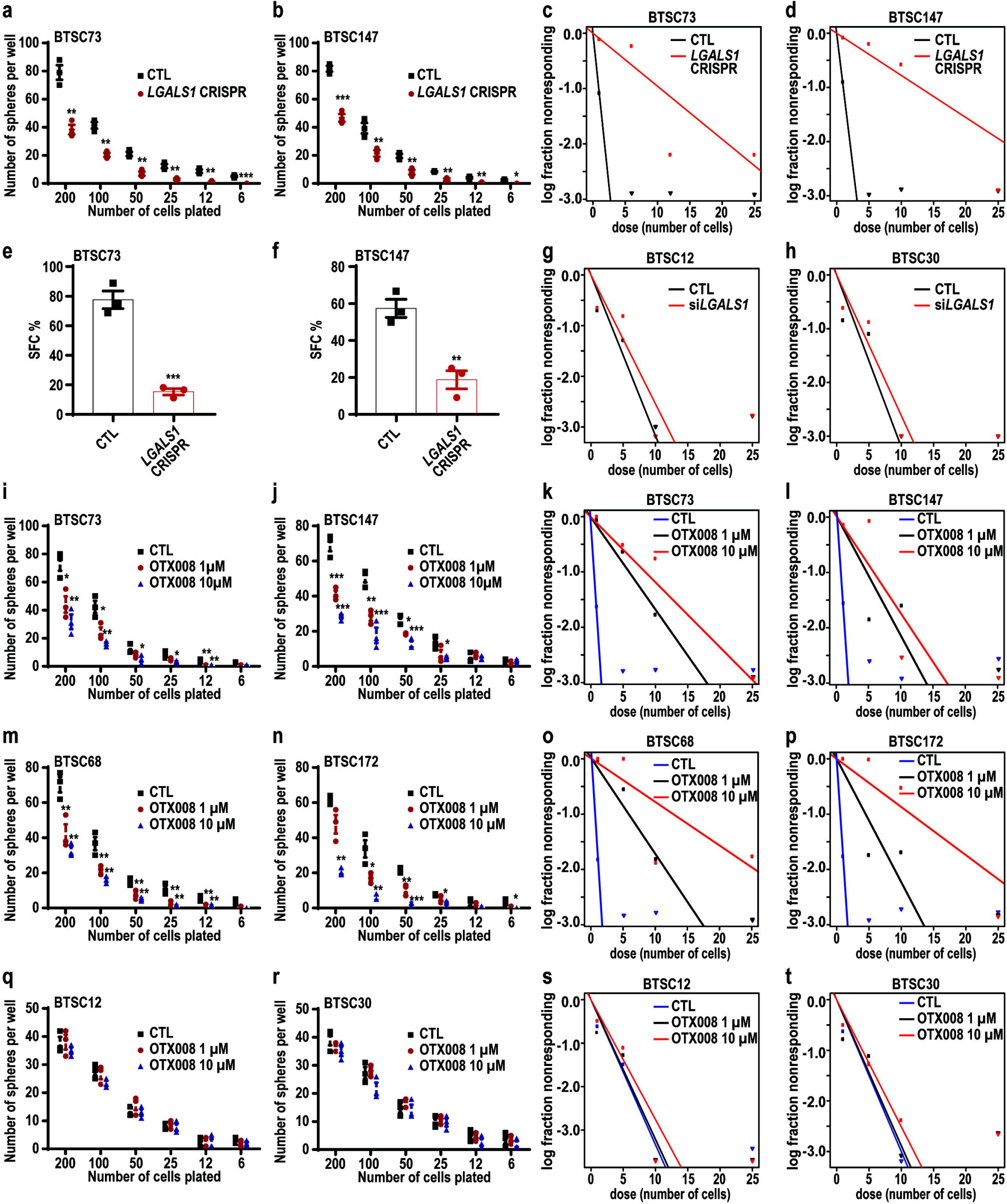
Galectin1 controls BTSC self-renewal. (**a-d**) *LGALS1* CRISPR and CTL EGFRvIII-expressing BTSCs were subjected to LDA (a-b) or ELDA (c-d). (**e-f**) EGFRvIII-expressing *LGALS1* CRISPR and CTL BTSCs were subjected to clonogenicity assay performed by culturing one single cell per well. (**g-h**) BTSCs that don’t harbour the EGFRvIII mutation were electroporated with siCTL or si*LGALS1* and subjected for ELDA analysis. (**i-p**) EGFRvIII-expressing BTSCs were subjected to LDA (i, j, m and n) or ELDA (k, l, o and p) following the treatment with 1 or 10 µM OTX008. (**q-t**) BTSCs that don’t harbour the EGFRvIII mutation were subjected to LDA (q-r) or ELDA (s-t) following the treatment with 1 or 10 µM OTX008. *p < 0.05, **p < 0.01, ***p < 0.001; unpaired two-tailed *t*-test (a, b, e and f); one-way ANOVA followed by Dunnett’s test (i, j, m and n), n = 3. Data are presented as the mean□±□SEM. See also Figure S6.

In another set of independent experiments, we sought to examine how EGFRvIII-expressing BTSCs or BTSCs lacking the mutation respond to the pharmacological inhibitor, OTX008. EGFRvIII-expressing BTSC lines (#73, 147, 68, 112, and 172) and BTSC lines that do not harbour EGFRvIII mutation (#12 and 30) were subjected to LDA and ELDA, following treatment with different concentrations of OTX008 or a vehicle control. Our analysis revealed that OTX008 resulted in a significant decrease in the number of spheres and self-renewal capacity in all the EGFRvIII-expressing BTSCs examined (**Figure 5i-p and Figure S6g-h**). Importantly, we observed a dose-dependent effect in OTX008-mediated suppression of self-renewal. In contrast, OTX008 had no significant impact on the self-renewal of BTSCs that do not harbour EGFRvIII mutation (**Figure 5q-t**). In summary, using genetic and pharmacological approaches, we establish a role of *LGALS1* in regulation of self-renewal of EGFRvIII-expressing BTSCs.

Notch signalling pathway plays vital roles in controlling glioma stem cells self-renewal (El-Sehemy et al., 2020; Park et al., 2017; Rajakulendran et al., 2019). Additional analysis of our RNA-seq data, revealed a panel of downregulated genes related to Notch signalling, including the Notch ligands, *JAG2* and *DLL1* in the *LGALS1* CRISPR BTSCs (**Figure S6i**). This raised the question of whether galectin1 controls Notch signalling in BTSCs via regulating the expression of ligands *JAG2* and *DLL1*. We performed immunoblotting analysis and observed a marked decrease in Jagged2 and DLL1 protein levels in *LGALS1* CRISPR in multiple EGFRvIII-expressing BTSCs (**Figure S6j-m**). Activation of the Notch pathway requires that the ligands bind the receptor resulting in the cleavage and release of the Notch intracellular domain (Kopan and Ilagan, 2009). We, thus, assessed the active Notch1 (cleaved Notch1) protein levels by WB and found a robust reduction in the cleaved Notch1 levels in *LGALS1* CRISPR BTSC73 and 147 compared to the corresponding controls (**Figure S6j-m**). Altogether, these data suggest that *LGALS1* may impact BTSC self-renewal via positive regulation of Notch signalling pathway.

### Pharmacological targeting of galectin1 sensitizes glioblastoma to IR and expands lifespan

Tumours that are highly resistant to IR including glioma, melanoma, and prostate cancer are found to express high levels of galectin1 (Navarro et al., 2020), raising the question of whether galectin1 may contribute to BTSC resistance in response to IR. To address this question, we employed ELDA analysis to assess the response of *LGALS1* CRISPR and control BTSC73 to 4 Gy of IR. IR induced 2.7-fold decrease in stem cell frequency of control BTSC73. Strikingly, deletion of *LGALS1* sensitized the BTSC73 response to IR by 14 folds (**Figure 6a**). Since exposure to IR provokes DNA damage that trigger cell death (Afshar et al., 2006), we analyzed cell death by annexin V and PI double staining and we found that loss of *LGALS1* resulted in a significant increase in IR-induced apoptosis, whereby 53% of irradiated *LGALS1* CRISPR BTSCs were positive for annexin V compared to only 22% in the irradiated control BTSCs (**Figure 6b-c**). These data suggest that *LGALS1* targeting sensitizes BTSCs to IR-induced cell death.

**Figure 6.**
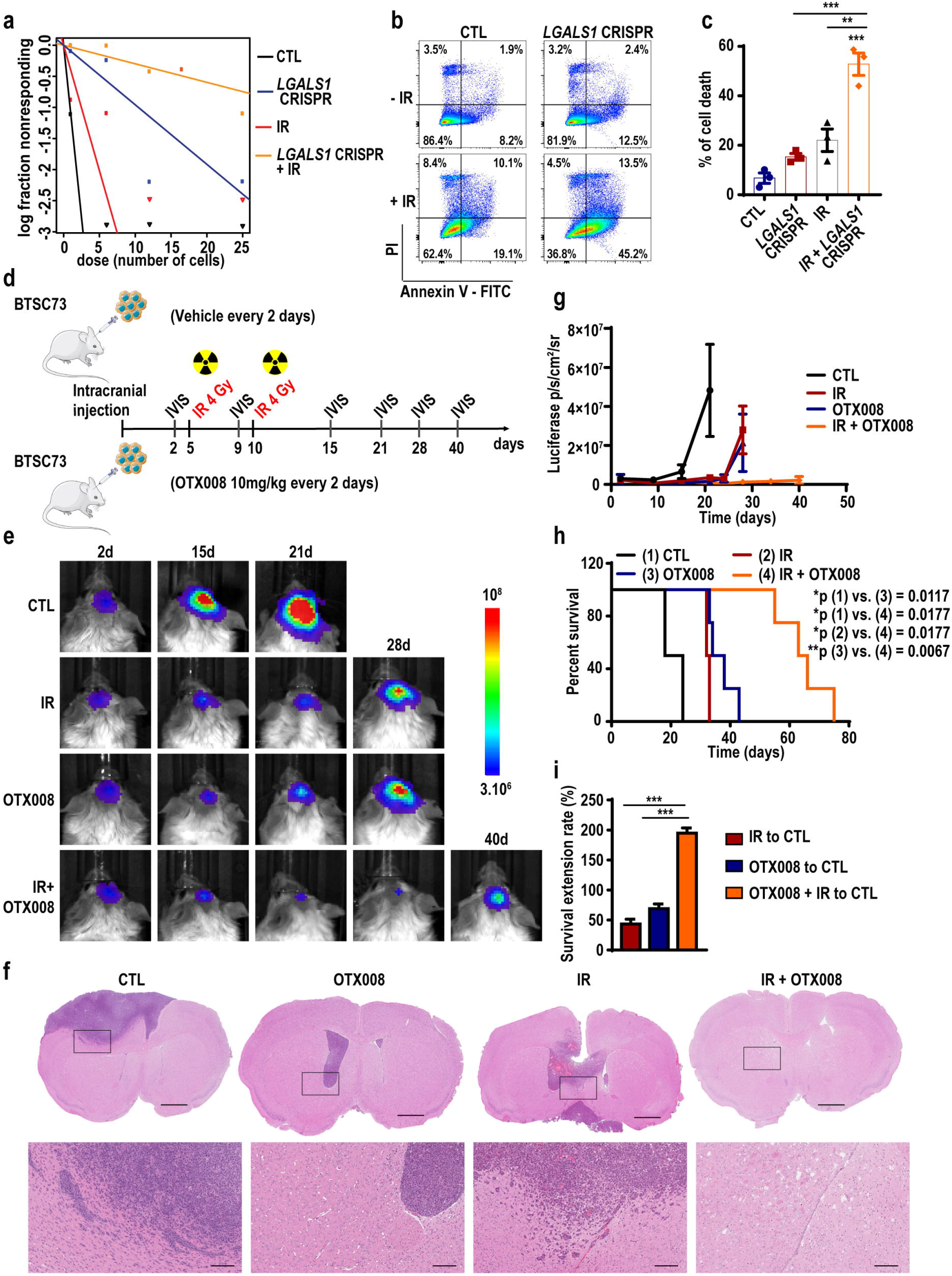
Pharmacological inhibition of galectin1 improves the response of patientderived brain tumours to IR. (**a**) ELDA was performed following 4 Gy of IR in *LGALS1* CRISPR or CTL BTSCs. (**b-c**) *LGALS1* CRISPR and CTL BTSC73 were subjected to IR (8□Gy). Apoptosis analysis was performed by flow cytometry 48□h following IR using annexin V and PI double staining. Representative scatter plots of flow cytometry analyses are shown (b). The percentage of cell death (annexin V positive cells) is presented in the histogram (c), n□=□3. (**d**) Schematic diagram of the experimental procedure is shown. BTSC73 were intracranially injected into SCID mice and then treated with OTX008, 4□Gy of IR or a combination of OTX008 and IR. (**e**) Representative bioluminescence real-time images tracing tumour growth are shown, n□=□6 mice. (**f**) Coronal sections of mouse brains were stained with hematoxylin and eosin on day 22 after injection. Representative images of 3 different tumour sections are shown. Scale bar = 1□mm, scale bar (inset) = 0.2 mm. (**g**) Intensities of luciferase signal were quantified at different time points, n = 6 mice. (**h**) KM survival plot was graphed to assess animal lifespan, n□=□6 mice. (**i**) Survival extension of mice bearing BTSC-derived tumours treated with OTX008, IR, or OTX008 + IR relative to those treated with the vehicle control. Data are presented as the mean□±□SEM. One-way ANOVA followed by Tukey’s test (c and i); log-rank test (h), *p < 0.05, **p < 0.01, ***p < 0.001.

To investigate the functional relevance of these findings to glioblastoma *in vivo*, we transplanted EGFRvIII-expressing BTSC73 into the brains of immunodeficient mice and assessed whether combinational therapy with IR and pharmacological inhibition of galectin1 by OTX008 improves the lifespan of the animals. Mice bearing BTSCs were treated with: vehicle control (PBS), OTX008 (10 mg/kg), IR (4 Gy) or the combination of OTX008 and IR (**Figure 6d**). At 21 days following surgery, mice receiving control BTSC73 formed invasive brain tumours and were at endpoint as assessed by major weight loss and neurological signs (**Figure 6e-h**). Exposure to IR alone delayed tumorigenesis whereby the mice were at endpoint at 30 days (**Figure 6e-i**). Treatment with OTX008 delayed tumourigenesis, inhibited invasiveness and extended lifespan by 40 days (**Figure 6e-i**). Strikingly, the combined OTX008 and IR significantly suppressed tumourigenesis as revealed by analysis of tumour volume and extended survival past 65 days (**Figure 6e-i**). Our data suggest that combination therapy with OTX008 and IR may be a promising avenue to overcome the resistance of EGFRvIII-expressing BTSCs and glioblastoma tumours.

### Galectin1 physically and functionally interacts with the TF HOXA5

To investigate the mechanisms by which *LGALS1* regulates BTSC, *LGALS1*-differentially regulated genes (**Figure S5**) were subjected to enrichment analysis of TF binding motifs using oPOSSUM-3 software to screen for TFs that could cooperate with galectin1 to reprogram BTSC transcriptional landscape. This analysis led to the identification of over-represented TF binding sites in the promoter of *LGALS1*-differentially expressed genes. The five TFs HOXA5, Pdx1, SRY, Nkx2-5, and ARID3A binding sites were significantly enriched within the promoter regions (Z-Score > mean + 2sd) (**Figure 7a**). HOXA5, a member of the homeobox TF family, was selected for further investigation based on the following criteria: **first**, among the five top identified TFs, HOXA5 ranked first whereby 70% of galectin1 potential target genes harboured the consensus motif for HOXA5 binding (**Figure 7a-b**). **Second,** we analyzed the protein expression levels of HOXA5, Pdx1, SRY, Nkx2-5, and ARID3A by immunoblotting in different patient-derived BTSCs and performed Pearson correlation analysis relative to galectin1 expression. We found that only the protein levels of HOXA5 and SRY correlates with galectin1 expression in BTSCs (**Figure 7c-d**, **Figure S7a-e**). **Third,** we employed TCGA public database and performed patient survival analysis. KM survival plots of glioblastoma patients were generated after clustering them based on the mRNA expression levels of *LGALS1* and each of the five predicted TFs. Our analyses demonstrated that patients with low expression of *LGALS1* and *HOXA5* had the best prognosis (**Figure 7e**), while the remaining four TFs had no significant impact on glioblastoma patient survival (**Figure S7f-i**)**. Fourth**, given that the observed phenotype of deregulated cell cycle progression and self-renewal in *LGALS1* CRISPR is associated with *LGALS1*-downregulated genes, we conducted further analysis in which only the *LGALS1*-downregulated genes were subjected to oPOSSUM-3 TF enrichment analysis. We found that HOXA5 was the only TF that had a Z-score above the threshold (Z-score > mean + 2SD) (**Figure S7j**). Furthermore, HOXA5 was recently shown to promote cellular proliferation and radioresistance in glioblastoma (Cimino et al., 2018). These analyses led us to examine if *LGALS1* cooperates with HOXA5 to alter BTSC transcriptional program. To begin with, we sought to validate the enrichment of HOXA5 binding sites on the *LGALS1-*differentially regulated genes. We queried the public available ChIP-seq data of human carcinoma cells (Yan et al., 2013) and analyzed the genomic distribution of HOXA5 ChIP-seq peaks relative to the *LGALS1*-differentially regulated genes in BTSCs. We found a significant abundance of HOXA5 ChIP-seq peaks in the 5′ untranslated and the promoter regions of *LGALS1* candidate target genes (**Figure 7f**), suggesting that *LGALS1*-differentially regulated genes are direct HOXA5 targets.

**Figure 7.**
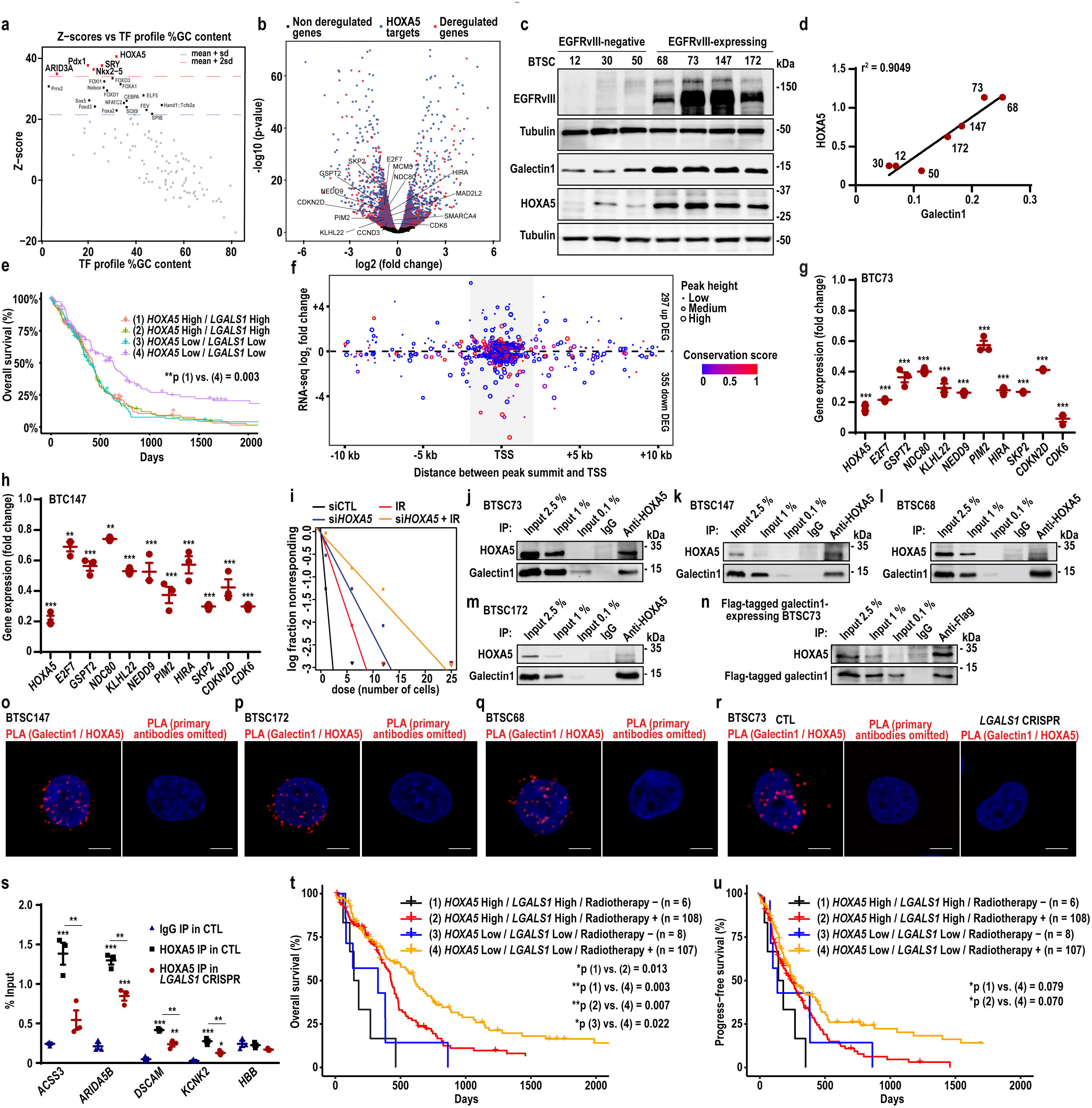
Galectin1 interacts with HOXA5 endogenously in patient-derived BTSCs. (**a**) *LGALS1*-differentially regulated genes were subjected to enrichment analysis of TF binding motifs using oPOSSUM-3 software. (**b**) Volcano plot representing the HOXA5 target genes among the *LGALS1*-differentially-regulated genes is shown. (**c**) BTSCs were analyzed by immunoblotting using the antibodies indicated on the blots. (**d**) Pearson correlation analysis of HOXA5 and galectin1 protein expression is shown. (**e**) KM survival plot describing the association between *LGALS1* and *HOXA5* expression and the survival of glioblastoma patients is shown. (**f**) Relative positions of HOXA5 ChIP-seq peaks to the adjacent TSS of *LGALS1*-differentially regulated genes are shown. The x-axis indicates the distance between peak centers and the TSS of adjacent *LGALS1*-differentially regulated genes. The y-axis denotes the expression ratios (log2) of the *LGALS1*-differentially regulated gene. Circle size indicates HOXA5 peak height, and color denotes the conservation score of HOXA5 peaks. (**g-h**) *HOXA5* KD (si*HOXA5*) and siCTL BTSCs were subjected to RT-qPCR analysis. (**i**) ELDA was performed following 4LGy of IR in si*HOXA5* vs. siCTL. (**j**-**m**) Endogenous Co-IP experiments were performed in different BTSC lines using an anti-HOXA5 antibody, followed by immunoblotting with galectin1 and HOXA5 antibodies. (**n**) Co-IP experiment was performed using anti-FLAG antibody, followed by immunoblotting with anti-FLAG and anti-HOXA5 antibodies. (**o**-**r**) PLA of galectin1 and HOXA5 were performed in different BTSC lines. Primary antibodies were omitted for the controls. Nuclei were stained with DAPI. Scale bar = 10 μm. (**s**) *LGALS1* CRISPR and CTL BTSC73 were subjected to ChIP using an antibody to HOXA5 followed by qPCR for HOXA5 candidate target genes. HBB locus was used as a negative control. (**t-u**) KM survival plot describing the association between *LGALS1* and *HOXA5* expression and the survival of glioblastoma patients treated with radiotherapy (microarray G4502A Agilent, level 3, n = 489). Data are presented as the meanL±LSEM, n = 3. Log-rank test (e, t and u); one-way ANOVA followed by Dunnett’s test (g and h); unpaired two-tailed *t*-test (s). *p < 0.05, **p < 0.01, ***p < 0.001. See also Figure S7.

Next, we induced KD of *HOXA5* by a pool of siRNA followed by RT-q-PCR for select cell cycle related genes that are downregulated in *LGALS1* CRISPR and possess HOXA5 binding motifs. Strikingly, similar to the effect of *LGALS1* deletion, we found a significant reduction in the mRNA levels of the 10 randomly selected cell cycle related genes in two patient-derived EGFRvIII-expressing BTSCs (#73 and 147) (**Figure 7g-h**). To establish if loss of *HOXA5* phenocopies the *LGALS1*-KO phenotype, we examined the impact of *HOXA5* KD on the self-renewal and radioresistance of EGFRvIII-expressing BTSCs as well as deregulation of Notch signalling. ELDA analysis revealed that similar to *LGALS1* deletion, KD of *HOXA5* significantly reduced stem cell frequency in BTSC73. Importantly, we observed a significant decrease in sphere formation frequency of irradiated BTSC upon KD of *HOXA5* (**Figure 7i**). In addition, similar to *LGALS1* deletion, KD of *HOXA5* attenuated the expression levels of ligands Jagged2 and DLL1, and reduced the active (cleaved) Notch1 levels (**Figure S7k-m**). These results established that *HOXA5* KD in BTSCs phenocopies the effects of *LGALS1* deletion and suggest a cross-talk between galectin1 and HOXA5 in reprogramming the BTSC transcriptional network.

Our findings in which we established a functional interaction between galectin1 and HOXA5 in regulation of BTSC, led us next to assess whether galectin1 physically interacts with HOXA5 in EGFRvIII-expressing BTSCs. We employed immunostaining and co-immunoprecipitation (Co-IP) experiments to examine protein-protein interactions. Strikingly, we found that HOXA5 physically interacts with galectin1 endogenously in different patient-derived EGFRvIII-expressing BTSCs (**Figure 7j-m**). In a follow up experiment, we expressed a FLAG-tagged galectin1 in BTSC73 and subjected the cells to immunoprecipitation using a FLAG antibody. Similar to endogenous Co-IP experiments, HOXA5 was detected in the co-immunoprecipitation complex (**Figure 7n**). To validate HOXA5-galectin1 interaction *in situ*, we performed proximity ligation assay (PLA). We detected significant PLA signals in multiple EGFRvIII-expressing BTSCs in which antibodies to HOXA5 and galectin1 were employed (**Figure 7o-r**). Importantly, deletion of *LGALS1* abolished the PLA interaction signal (**Figure 7r**). TFs bind to the gene regulatory regions to control gene expression. We therefore examined if galectin1 is required for HOXA5 TF activity by promoting its DNA binding activity. We performed ChIP assay for HOXA5 in *LGALS1* CRISPR and CTL BTSC73, followed by qPCR for select *LGALS1*-downregulated genes that possess HOXA5 binding motifs. ChIP-qPCR revealed an enrichment of HOXA5 on the promoter regions of the potential target genes in the BTSC73. Strikingly, silencing of *LGALS1* significantly impaired the binding of HOXA5 to the promoter of its target genes (**Figure 7s**). Together, these data established that galectin1 forms a complex with HOXA5 and this interaction is essential for HOXA5 TF activity.

Finally, having established that galectin1 mediates its effects through a cross-talk with HOXA5, we sought to determine the prognostic value of *LGALS1*/*HOXA5* expression in human glioblastoma patients in response to IR. We employed the TCGA (microarray G4502A) dataset in which IR treatment and patient’s response to therapy were available. We generated KM survival plots for glioblastoma patients after clustering them based on the mRNA expression levels of *LGALS1* and *HOXA5* into high (above median) and low (below median). Our analyses demonstrated that patients with low expression of *LGALS1* and *HOXA5* had best prognosis in response to IR (**Figure 7t-u**). Our data highlights the importance of galectin1*/*HOXA5 targeting in combination with IR as a potential promising therapeutic regimen for glioblastoma patients.

## Discussion

In the present study, we report a role for the *LGALS1* gene, encoding the carbohydrate binding protein, galectin1, in maintaining the EGFRvIII/STAT3 oncogenic signalling in glioma stem cells and conferring glioblastoma resistance to therapy. Beginning with loss and gain of function studies using genetic and pharmacological approaches we found that *LGALS1* is upregulated in an EGFRvIII and STAT3 dependent manner. We employed ChIP and luciferase reporter assays and showed that STAT3 directly occupies the promoter of *LGALS1* in EGFRvIII-expressing BTSCs and upregulates its expression. Importantly, using genetic and pharmacological approaches we established that inhibition of this signalling pathway robustly impairs the self-renewal and growth of BTSCs and sensitizes the response of glioblastoma tumours to IR. Finally, via RNA-seq analysis followed by unbiased TF enrichment analysis, ChIP-seq analysis, PLA and endogenous Co-IP experiments, we established that galectin1 physically and functionally interacts HOXA5 to reprogram BTSC transcriptional network and confer glioblastoma resistance to IR. Our data unravel a STAT3/*LGALS1*/HOXA5 signalling axis that tightly regulates BTSC and promotes tumourigenesis of EGFRvIII subtype of glioblastoma.

Epigenomic and transcriptomic analyses have revealed that EGFRvIII specifically controls glioblastoma response to therapy via modulating transcription (Liu et al., 2015). EGFRvIII forms a complex with STAT3 to control gene regulation and glioblastoma tumourigenesis (de la Iglesia et al., 2008; Puram et al., 2012). Our findings that *LGALS1* is a direct transcriptional target of EGFRvIII/STAT3 and promotes tumourigenesis of EGFRvIII subtype of glioblastoma, raises important implications for developing novel therapeutic regimen that includes a combination of EGFRvIII/STAT3 and galectin1 inhibitors.

Aberrant activation of STAT3, plays an important role in the mesenchymal subtype of glioblastoma (Carro et al., 2010; Hara et al., 2021; Verhaak et al., 2010). In a recent study, the cytokine OSM, produced by macrophages, was shown to interact with cancer cells to activate STAT3 and induce the transition of glioblastoma cells into mesenchymal-like state (Hara et al., 2021). Importantly, our analysis revealed that OSM mediated phosphorylation of STAT3 resulted in increased expression of *LGALS1*. Most importantly, deletion of *LGALS1* resulted in downregulation of the mesenchymal gene signature. This finding holds promise to investigate the therapeutic potential of targeting *LGALS1* in mesenchymal subtype of glioblastoma.

Notch signaling plays important roles in regulation of glioma stem cell fate (El-Sehemy et al., 2020; Park et al., 2017; Rajakulendran et al., 2019; Ranganathan et al., 2011). Blocking of Notch1 or its ligands, DLL1 and Jagged1, has been shown to inhibit the proliferation of glioma cells (Purow et al., 2005). Furthermore, aberrant expression of various Notch components correlates significantly with aggressive high grade glioma and poor prognosis (Dell’albani et al., 2014; Phillips et al., 2006; Somasundaram et al., 2005; Zhang et al., 2017). In our studies, we found that loss of either *LGALS1* or *HOXA5* significantly attenuates the expression of the ligands DLL1 and Jagged2 and the cleavage of Notch1. Our data suggests that galectin1/HOXA5 complex may cooperate with the Notch signaling to regulate BTSC self-renewal. In support of this model, *LGALS1* is shown to promote lung cancer progression by activation of the Jagged2/Notch1 signalling pathway (Hsu et al., 2013).

Using murine astrocytes, STAT3 has been shown to play opposing roles in glial transformation depending on the mutational profile of the tumour. Although, STAT3 forms a complex with the EGFRvIII in the nucleus to mediate EGFRvIII-induced glial transformation. STAT3 plays a tumour suppressive role in the PTEN deficient tumours (de la Iglesia et al., 2008). Similarly, HOXA5 has been shown to play opposing roles in tumourigenesis. For example, HOXA5 is a positive regulator of TP53 in solid tumours such as breast and lung cancer (Raman et al., 2000). In contrast, HOXA5 provides a selective advantage for gain of chromosome 7 in IDH wild-type glioblastoma and a more aggressive glioma phenotype (Cimino et al., 2018). We provide experimental evidence that similar to STAT3, HOXA5 functions in an oncogenic capacity in the genetic background of EGFRvIII. Furthermore, STAT3 and HOXA5 function on the same signalling pathway to regulate galectin1 expression and function. Although the mechanisms that regulate HOXA5 function remain to be investigated, our data suggests that loss of galectin1 impairs the binding of HOXA5 to DNA and its TF activity. Whether galectin1 functions as a co-transcriptional regulator in this context remains to be investigated in future studies.

In conclusion, we report a role for the carbohydrate binding protein, galectin1, in regulation of glioma stem cells and glioblastoma progression via a cross talk with the TF HOXA5. Our data highlights the importance of galectin1-HOXA5 targeting as a promising approach to deplete malignant cancer stem cells and suppress glioblastoma tumourigenesis.

## Acknowledgements

This work was supported by grant (# 25139) from Cancer Research Society and grant (#440097) from the Canadian Institute of Health Research. AJA is a Fonds de la recherche en santé du Québec (FRQS) investigator. AB is supported by an FRQS postdoctoral fellowship. We thank Dr. Samual Weiss at the University of Calgary for sharing BTSC12, 30, 50, 68, 73,147 and Dr. Keith Ligon at Harvard Medical School for the generation of BTSC112 and 172. We thank Mehdi Haghi for technical assistance. We thank staff at the Lady Davis Institute Animal Core Facility and Christian Young at Fluorescence-Activated Cell Sorting (FACS) facility at the Lady Davis Institute for Medical Research – Jewish General Hospital for their help with our studies.

## Author Contributions

Performed experiments: A.S., A.B., I.F. and A.J.-A.; Designed experiments and analyzed data: A.S., A.B., I.F., V.D.S. and A.J.-A.; bioinformatics analyses of RNA-seq and TCGA data: A.S., A.H.-C., A.M. and H.S.N.; Wrote the paper: A.S. and A.J.-A.; Conceived the research program and provided funding and mentorship: A.J.-A.

## RESOURCE AVAILABILITY

### Lead contact

Further information and requests for resources and reagents should be directed to and will be fulfilled by the lead contact, Arezu Jahani-Asl.

### Materials availability

This study did not generate new unique reagents.

### Data and code availability

RNA-seq data generated in this study are available in GEO database under the accession number GEO: GSE180981.

## EXPERIMENTAL MODEL AND SUBJECT DETAILS

### Animals

All animal experiments were conducted under the institutional guidelines and were approved by McGill University Animal Care Committee (UACC). 8-week-old male SCID mice (Charles River) were used and randomly assigned. Housing room temperature and relative humidity were adjusted to 22.0L±L2.0L°C and 55.0L±L10.0%, respectively. The light/dark cycle was adjusted to 12 h lights-on and 12 h lights-off. Autoclaved water and irradiated food pellets (Tecklad, #2918) were given ad libitum.

### Cell lines

The human BTSC lines 112 and 172 were generously provided by Dr. Keith Ligon at Harvard Medical School. BTSC lines were generated following surgery with informed consent of adult glioblastoma patients following the BWH/Partners IRB protocol for use of excess/discarded tissue at Harvard University. BTSCs 12, 30, 50, 68, 73, and 147 were provided by Dr. Samuel Weiss at the University of Calgary (Chesnelong et al., 2019; Cusulin et al., 2015). Cells were characterized for major mutations in glioblastoma including EGFRvIII, p53, PTEN, and IDH1 status (Jahani-Asl et al., 2016). BTSCs 68, 73, 147, and 172 that naturally harbour EGFRvIII mutations, and BTSCs 12, 30 and 50 that do not harbour the mutation, were used in this study. Prior to use, BTSCs were recovered from cryopreservation in 10% dimethyl sulfoxide and cultured in Nunc ultra-low attachment flasks as maintained in neurospheres in serum free NeuroCult NS-A medium (Stemcell Technologies, #05750) supplemented with 100 U/mL penicillin, 100 µg/mL streptomycin (Sigma Aldrich, #P4333), heparin (2 μg/mL, Stemcell Technologies, #07980), human EGF (20 ng/mL, Miltenyi Biotec, #130-093-825), and human FGF (10 ng/mL, Miltenyi Biotec, #130-093-838). BTSCs were passaged up to 7 times and all cell lines were tested negative for mycoplasma. BTSCs were from glioblastoma patients of the following age and sex: BTSC12 (RRID: CVCL_RU28), 59 years old male; BTSC30 (RRID: CVCL_UK45), 67 years old male; BTSC50 (RRID: CVCL_UK38), 61 years old male; BTSC68 (RRID: CVCL_UK52), 59 years old male; BTSC73 (RRID: CVCL_UK54), 52 years old male; BTSC147 (RRID: CVCL_UK63), 55 years old male. For BTSC112 (RRID: CVCL_WW37) and BTSC172 (RRID: CVCL_IP12) sex and ages are not available.

Mouse astrocytes were cultures in DMEM supplemented with 10% fetal bovine serum, 1 mM L-Glutamine, 100 U/mL penicillin, and 100 μg/mL streptomycin (de la Iglesia et al., 2008). All cells were kept in a humidified 5% CO_2_ incubator at 37°C.

## METHOD DETAILS

### Generation of transgenic BTSCs

We employed 3 different approaches to delete *LGALS1* and *HOXA5* in patient-derived human BTSCs. First, genetic deletion of *LGALS1* and *HOXA5* was achieved using CRISPR. Briefly, two gRNAs were designed using off-spotter software to delete exon 2-4 resulting in a 2.673 kb deletion of *LGALS1* gene. To delete *HOXA5*, gRNAs were designed to delete exon 1-2 resulting in a 2.218 kb deletion of *HOXA5* gene. gRNA-1 and -2 were cloned into pL-CRISPR.EFS.GFP (Addgene, #57818) and pL-CRISPR.EFS.tRFP (Addgene plasmid #57819), respectively (Heckl et al., 2014). 5 ug of each construct were nucleofected into BTSC73 or BTSC147 using an AMAXA nucleofector 2b device (Lonza, #AAB-1001). The GFP and RFP positive cells were then sorted two days post-electroporation and plated clonally using FACSAria Fusion. Genomic DNA was isolated from each clone and screened for *LGALS1* or *HOXA5* deletion via PCR using specific internal and external primers around the site of the deletion. This led to the identification of monoallelic deletion, biallelic deletion and non-deletion clones. *LGALS1* mRNA and protein levels were analyzed by RT-qPCR and WB, respectively, to assess KD levels (**Figure S3a-e**). gRNAs and screening primers used for CRISPR/Cas9 system are shown in Table S1.

Second, the transgenic *LGALS1* KD BTSCs were generated via lentivirus carrying two different *LGALS1* shRNA plasmids (OriGene, #TL311756). *LGALS1* KD BTSC73 lines were established by antibiotic selection (0.5 μg/mL puromycin). As control, a lentivirus carrying a non-targeting construct was used.

Third, we conducted transient KD of *LGALS1* and *HOXA5* using siRNA approach. ON TARGET-plus SMART pool human *LGALS1* siRNA (Dharmacon, #L-011718-00-0005), human *HOXA5* siRNA (Dharmacon, #L-017574-00-0005) and ON TARGET-plus non-targeting pool (Dharmacon, #D-001810-10-05) were used. siRNA (100 nM) were nucleofected into BTSCs (10^6^ cells) and cultured in BTSC media at 37°C in a humidified atmosphere of 5% CO_2_. Transient KD of *STAT3* and *EGFR*/*EGFRvIII* was achieved using siRNA approach. ON TARGET-plus SMART pool human *STAT3* siRNA (Dharmacon, #L-003544-00-0005) and *EGFR*/*EGFRvIII* siRNA (Dharmacon, #L-003114-00-0005) were used. siRNA (100 nM) were nucleofected into BTSCs (10^6^ cells) and cultured in BTSC media at 37°C in a humidified atmosphere of 5% CO_2_.

To generate FLAG-tagged galectin1 expressing BTSCs, BTSCs were electroporated with pCMV6-*LGALS1*-Myc-DDK mammalian vector (OriGene, #RC204674).

### Inhibitors

Phosphorylation of EGFR and EGFRvIII were inhibited by lapatinib (Sigma, #CDS022971). Phospho-STAT3 was inhibited using WP1066 (Sigma, #573097) and S3I-201 (Sigma, #SML0330). OTX008 (MedChemExpress, #HY-19756) was used to inhibit galectin1.

### Ionizing radiation

For ELDA and measurement of cell death, BTSCs were dissociated to single cell suspension using Accumax (Innovative Cell Technologies, #AM105). BTSCs were plated and irradiated with either 4 or 8□ Gy using the X-Ray Irradiation System (Faxitron MultiRad 225).

### OTX008 treatment

For LDA, ELDA and measurement of cell death, BTSCs were dissociated to single cell suspension, plated and treated with different OTX008 concentrations or vehicle for 7 days.

### LDA, ELDA, and clonogenic assay

For LDA, BTSCs were dissociated into single cell suspension using Accumax, counted and plated in 96-well plate at different densities ranging from 200 to 6 cells per well in triplicates. Spheres were counted 7 days after plating.

For ELDA experiments, decreasing numbers of BTSCs per well (dose: 25, 12, 6, 3 and 1) were plated in a 96-well plate with a minimum of 12 wells/dose. Seven days after plating, the presence of spheres in each well was recorded and analysis was performed using the software available at http://bioinf.wehi.edu.au/software/elda/ (Hu and Smyth, 2009). For clonogenic assays, BTSCs were seeded in 96-well plates at clonal density of 1 cell per well. Seven days after plating, the presence of neurospheres in each well was recorded and the sphere forming capacity was calculated as a percentage of positive wells (Eyler et al., 2011).

### Cell population growth assay

BTSCs were dissociated into single cell suspension using Accumax, counted and plated at a density of 2 x 10^4^ cells per well. Following 24-, 48-, 72- and 96 h of plating, live cells were counted by trypan blue exclusion with Countess™ II FL Automated Cell Counter.

### EdU proliferation assay

BTSCs were dissociated into single cell suspension using Accumax, counted and plated at a density of 10^6^ cells per well. Cells were incubated with 10 µM EdU at the time of plating. After 22h, BTSCs were fixed, permeabilized and stained using the Click-iT™ EdU cell proliferation kit (Thermo Fisher Scientific, #C10337) according to the manufacturer’s protocol. Fluorescence was analyzed by flow cytometry (BD FACS CantoII) and images were acquired using a 10X objective on an Olympus IX83 microscope with an X-Cite 120 LED from Lumen Dynamics and an Olympus DP80 camera. Data were analyzed using the FlowJo software. The proportion of cells that incorporated EdU was determined as the ratio of EdU positive cells to the total number of cells.

### Cell cycle analysis

BTSCs were dissociated into single cell suspension using Accumax and 10^6^ cells were plated. After 22h, cells were dissociated to single cell suspension, harvested and fixed with 70% of ethanol overnight at 4°C. The cells were washed with 1X PBS and stained with FxCycle™ PI/RNase staining solution (Thermo Fisher Scientific, #F10797). The fluorescence was analyzed by flow cytometry (BD FACS CantoII). The fraction of G0/G1, S and G2 phase cells was calculated using the FlowJo software.

### Cell proliferation and cell death assessment

BTSCs were dissociated to single cell suspension using Accumax and seeded at a density of 200 cells/well, in a 96-well plate. Cell proliferation was evaluated 7 days post-plating using CellTiter-Glo® Luminescent Assay (Promega, #G7570) according to the manufacturer’s protocol, as previously described (Liu et al., 2020). Briefly, CellTiter-Glo reagent was added to the culture media. After 2 min under orbital agitation, luminescence was recorded. The luminescent signal is proportional to the amount of ATP present and directly proportional to the number of cells present in culture.

BTSCs cell death was evaluated 48 h following IR and 7 days following OTX008 treatment. Cell death was determined using TACS annexin V-FITC apoptosis detection kit (R&D systems, #4830-01-K) according to the manufacturer’s protocol. Briefly, 10^6^ cells were seeded in a 25 cm^2^ flask. Co-staining with TACS annexin V-FITC and PI was performed on single cell suspension following the manufacturer’s instructions. The fluorescence was analyzed by flow cytometry (BD FACS CantoII). Data were analyzed using the FlowJo software. Both early apoptotic (annexin V-positive, PI-negative) and late apoptotic (annexin V-positive and PI-positive) cells were included in the cell death plots.

### Collagen type I invasion assay

After 7 days of culture, BTSCs neurospheres were resuspended in 0.4 mg/ml of collagen I (Sigma-Aldrich, #C7661) in BTSC media without growth factors. 100 μL of the collagen and neurosphere mixture were plated in triplicates in a 96-well plate and incubated on ice for 5 min. Then, the plate was transferred to a 37 °C incubator for 5 min to allow collagen polymerization. Phase-contrast time-lapse images of the spheres were captured at different time-points using a 10x objective on an Olympus IX83 microscope. Invasion areas were measured after 24 h with Fiji software as the mean of three independent wells (Schindelin et al., 2012). An invasive index was calculated by taking the ratio of the sphere invasion areas after 24 h and the sphere surface at the time of plating.

### Protein immunoprecipitation

Immunoprecipitations were performed from whole cell lysate of BTSCs. Cells were lysed for 30 min on ice in lysis buffer (50 mM Tris pH 7.5, 150 mM NaCl, 2 mM MgCl_2_, 0.5 mM EDTA, 0.5% Triton X-100, protease inhibitor cocktail). Lysates were cleared by centrifugation (14,800 *g*, 20 min, 4°C) and subsequently incubated with anti-HOXA5 antibody (Abcam, #ab82645), anti-DDK (FLAG) antibody (OriGene, #TA150014) or rabbit IgG (Cell Signaling, #3900S) as a control. Primary antibody incubations were carried out overnight at 4°C, followed by a 1 h incubation at room temperature with Dynabeads Protein G magnetic beads (Thermo Fisher Scientific, #10003D). Beads were washed three times with lysis buffer and eluted by boiling in SDS sample buffer. Immunoprecipitates were analyzed by immunoblotting using the indicated antibodies.

### Immunoblotting and antibodies

Total proteins were harvested in RIPA lysis buffer containing protease and phosphatase inhibitors (Thermo Fisher Scientific, #A32959). Protein concentration was determined by Bradford assay (Bio-Rad), after which samples were subjected to SDS-PAGE and electroblotted onto Immobilon-P membrane (Millipore). Membranes were blocked in 5% bovine serum albumin in TBST, before sequential probing with primary antibodies and HRP-conjugated secondary antibodies in blocking solution. Target proteins were visualized by ECL (Biorad) using ChemiDoc Imaging System (Biorad). The following antibodies were used: galectin1 (1:1000, Cell Signaling, #12936), HOXA5 (1:500, Abcam, #ab82645), phospho-EGFR (Tyr1068) (1:1000, Abcam, #ab40815), EGFR (1:1000, Abcam, #ab32077), EGFRvIII (1:500, (Lavictoire et al., 2003)), α-tubulin (1:5000, Abcam, #ab4074), GAPDH (1:1000, Cell Signaling, #2118), STAT3 (1:1000, Cell Signaling, #9139), phospho-STAT3 (Tyr705) (1:1000, Cell Signaling, #9138), DLL1 (1:300, Proteintech, #20230-1-AP), Notch1 (1:500, Proteintech, #20687-1-AP) and Jagged2 (1:500, Cell Signaling, #2210).

### Immunofluorescence

For immunostaining, BTSCs were plated on Lab-Tek II, CC2-treated chamber slide system (Thermo Fisher Scientific, #154941) in media containing 10% FBS, for 30□min. Cells were washed with PBS and fixed with 4% paraformaldehyde for 15□min at room temperature. Next, cells were permeabilized with 0.5% Triton X-100 (Sigma Aldrich, #T8787) for 20□min and blocked for 1□h with 5% normal donkey serum (NDS) in 1X-PBS. The cells were then incubated overnight at 4□°C with primary antibodies to galectin1 (1:100, Cell Signaling, #12936S) diluted in 5% NDS-1X PBS. Cells were washed with 1X PBS and then incubated with secondary Alexa fluor 488 goat anti-rabbit (1:500, Cell Signaling, #4412S) antibody for 1□h. 2□μg/mL DAPI (Thermo Fisher Scientific, #D1306) was used to detect the nuclei and ProLong Gold Antifade Mountant (Thermo Fisher Scientific, #P36934) was used for mounting. Images were captured using a 63X objective on a laser scanning confocal microscope (ZEISS LSM 800).

### Duolink proximity ligation assay

PLAs were performed using a Duolink In Situ Red Starter Kit (Sigma, #DUO92101) according to the manufacturer’s instructions. Briefly, BTSC were plated on Lab-Tek II, CC2-treated chamber slides in media containing 10% FBS, for 30□min. Cells were washed with PBS and fixed with 4% paraformaldehyde for 15□min at room temperature. Next, cells were permeabilized with 0.5% Triton X-100 for 20□min, blocked using Duolink blocking solution, and then incubated with primary antibodies at 4□°C overnight. After washing, the oligonucleotide (Minus and Plus)-conjugated secondary antibodies were added and incubated for 1□h at 37□°C. Subsequently, cells were washed and incubated with the ligation solution for 30□min at 37□°C. The ligated nucleotide circles were amplified using polymerase via the addition of the amplification solution followed by incubation for 100□min at 37□°C. The slides were washed briefly, and Duolink In Situ Mounting Medium with DAPI (DUO82040, Sigma) was added to each sample to stain the nuclei. The visualized fluorescence PLA signals were captured using a 63X objective on a laser scanning confocal microscope (ZEISS LSM 800).

### Whole-transcriptome analyses (RNA-seq)

Total RNAs were isolated from cells using TRIzol reagent (Invitrogen) according to the manufacturer’s instructions. The quality of RNA was assessed by bioanalyzer before sequencing. Libraries for poly(A)^+^ RNA were prepared according to the Illumina protocol. Libraries were sequenced on Illumina NextSeq 500 High Output Flow Cell. RNA-seq reads were mapped to the human genome assembly (hg38) using HISAT2 (Kim et al., 2015). The number of aligned reads per gene was obtained with HTSeq (Anders et al., 2015) and gene annotations were retrieved from GENCODE (Frankish et al., 2019). Genes with an average read counts smaller than 10 were filtered out. Differential expression analysis was performed with DESeq2, *p*-values were corrected by independent hypothesis weighting (Ignatiadis et al., 2016; Love et al., 2014). Genes with an adjusted *p*-value smaller than 0.1 were considered statistically differentially expressed. GSEA was done with the fgsea R package (Korotkevich et al., 2019). The Log fold changes from DESeq2 were used to create a pre-ranked gene-list. The Reactome database, gene sets (Neftel et al., 2019) and the Chemical and Genetic Perturbation Gene sets (c2.cgp.v6.0) from MSigDB were used as references for pathways (Fabregat et al., 2018; Liberzon et al., 2011).

### Gene expression analysis

Total RNAs were isolated from cells using TRIzol reagent (Invitrogen) according to the manufacturer’s instructions. RNAs were then subjected to reverse transcription using the 5X All-In-One RT MasterMix cDNA synthesis system (abm, #G492). RT-qPCR was performed using the fluorescent dye SYBR Green (Biorad, #1725271). mRNA expression levels were then normalized to the housekeeping gene beta-glucuronidase (GUSB). qPCR primers are shown in Table S1.

### Chromatin immunoprecipitation

Cells were washed with PBS containing protease inhibitors (Thermo Fisher Scientific, #A32959) before fixation. Cells were cross-linked with 1% formaldehyde in PBS for 10 min, and quenched with a solution containing 0.125 M glycine in PBS for 5 min at room temperature. Washing, fixing and quenching of the cells were performed in 15-ml Falcon tubes with cells rotating at room temperature. Following quenching, cells were washed twice with PBS containing protease inhibitors, and cell pellets were collected by spinning at 150 *g*. for 10 min at 4 °C. Pellets were dissolved in ChIP lysis buffer (40 mM Tris-HCl, pH 8.0, 1.0% Triton X-100, 4 mM EDTA and 300 mM NaCl) containing protease inhibitors. Chromatin was fragmented by sonication in a water bath Bioruptor at 4 °C to an average length of 500 base pairs (bp). The lysates were spun at 12,000 *g*. for 15 min, and the supernatant was diluted 1:1 in ChIP dilution buffer containing 40 mM Tris-HCl, pH 8.0, and 4 mM EDTA plus protease inhibitors. Immunoprecipitation was done using a ChIP-grade HOXA5 antibody (Abcam, # ab82645), total STAT3 antibody (Cell Signaling Technology, #9139), mouse IgG antibody (Cell Signaling, #5415S), or rabbit IgG antibody (Cell Signaling, #3900S). Antibody-protein-DNA complexes were collected, washed and eluted, and the cross-links were reversed as described previously (Soleimani et al., 2013). Immunoprecipitated DNA was analyzed by q-PCR and binding enrichment was expressed as % of input. Primer sequences are shown in Table S1.

### TF binding sites enrichment analysis

TF binding sites over-representation analysis was performed using oPOSSUM (v.3.0, Single Site Analysis tool) (Kwon et al., 2012). Differentially downregulated genes were used as targets and all genes measured by our RNA-seq were used as background (Conservation cutoff: 0.6; Matrix score threshold: 85%; upstream/downstream region: 5kb/5kb; JASPAR CORE Profiles: All vertebrate profiles).

### Patient survival analysis

Patient and expression data were retrieved from TCGA (Agilent, G4502A) (Colaprico et al., 2016). Patients were divided into high and low groups by expression of *HOXA5* and *LGALS1* (separated at the median) and if they were treated with radiotherapy (+/-). Overall and progress-free survival was analyzed by KM curves. Pairwise comparisons between groups, using a log-rank test, were corrected with Benjamini-Hochberg multiple testing procedure. Adjusted *p*-values smaller than 0.1 were considered statistically significant.

### ChIP-seq analysis

HOXA5 ChIP-seq reads from colon adenocarcinoma (GSE51142) were mapped to the human genome assembly (hg38) using Bowtie2 (Langmead and Salzberg, 2012; Yan et al., 2013). Peak calling was carried out using MACS (with thresholds p-value < 10^−5^ and FDR < 0.2) (Zhang et al., 2008). Averaged conservation scores for HOXA5 ChIP-seq peaks were calculated based on phastcon scores (0∼1; UCSC 30-way). Transcription start sites were obtained from bioMart (Smedley et al., 2015). Peak height was calculated as the number of tags per 100 bp. Distance between TSS and ChIP-seq summit was calculated using Bedtools (Quinlan, 2014).

### Dual-luciferase Reporter Assay

The upstream 376 bp region of the human *LGALS1* transcriptional start site was cloned into the pGL4.23 (Promega) vector to generate the *LGALS1* luciferase reporter gene (*LGALS1* pGL4.23) by digesting the plasmid and the annealed primer pair using EcoRV (NEB, #R0195L) and HindIII (NEB, #R0104L) and ligating them with T4 DNA ligase (NEB, #M0202L). Primer sequences are shown in Table S1.The construct was confirmed by DNA sequencing. BTSCs and mouse astrocytes were electroporated with the *LGALS1* pGL4.23 construct or the empty pGL4.23. Luciferase assays were performed 48 h after transfection with the Dual-Luciferase Reporter Assay system (Promega, #E1910) with a GloMax Luminometer (Promega). In all experiments, cells were electroporated with a Renilla firefly reporter control and the firefly luminescence signal was normalized to the Renilla luminescence signal.

### Stereotaxic injections and bioluminescent imaging

For intracranial injections, 3□×□105 luciferase-expressing BTSCs were stereotactically implanted into the right striata (0.8□mm lateral to the bregma, 1□mm dorsal and 2.5□mm from the pial surface) of randomized 8-week-old male SCID mice. Mice were randomly assigned to the treatment or vehicle. For OTX008 treatment, seven days post BTSCs injection, mice were administered intraperitoneally 10 mg/kg OTX008 or vehicle control every two days. For IR treatment, mice received 4□Gy of IR using the X-Ray Irradiation System (Faxitron MultiRad 225) five and ten days following BTSCs injections (Sharanek et al., 2020). To examine tumour volume, the animals were intraperitoneally injected with 200□□μL of 15□mg/mL D-luciferin (Thermo Fisher Scientific, #88292), anesthetized with isoflurane inhalation, and subjected to bioluminescence imaging with a CCD camera (IVIS, Xenogen) on a weekly basis. All bioluminescent data were collected and analyzed using IVIS software. For KM survival plots, mice were collected when they showed signs of tumour-related illness

### Subcutaneous xenografts

Luciferase-expressing BTSCs (10^6^ cells) were subcutaneously injected into flanks of 8-week-old male SCID mice. Mice were randomly assigned to the treatment or vehicle. For OTX008 treatment, seven days post BTSCs injection, mice were administered intraperitoneally 10 mg/kg OTX008 or vehicle control every two days. Tumour growth was evaluated by luciferase imaging. Mice were killed when mice developed neurological signs and ulcerated tumours, and the tumours were removed and weighted.

## QUANTIFICATION AND STATISTICAL ANALYSIS

Statistical analysis was performed using ANOVA and Student’s *t*-test, with the aid of GraphPad software 7. Two-tailed and unpaired t-tests were used to compare two conditions. One-way ANOVA with Tukey’s or Dunnett’s post hoc analyses were used for analyzing multiple groups. Data are shown as mean with standard error of mean (mean□±□SEM). The log-rank test was used for statistical analysis in the KM survival plot. p*-*values of equal or less than 0.05 were considered significant and were marked with an asterisk on the histograms. p*-*values of less than 0.05 are denoted by *, p*-*values of less than 0.01 are denoted by **, and p-values of less than 0.001 are denoted by *** on the histograms.

## References

Afshar, G., Jelluma, N., Yang, X., Basila, D., Arvold, N.D., Karlsson, A., Yount, G.L., Dansen, T.B., Koller, E., and Haas-Kogan, D.A. (2006). Radiation-induced caspase-8 mediates p53-independent apoptosis in glioma cells. Cancer Res 66, 4223–4232. 10.1158/0008-5472.CAN-05-1283.

Anders, S., Pyl, P.T., and Huber, W. (2015). HTSeq--a Python framework to work with high-throughput sequencing data. Bioinformatics 31, 166–169. 10.1093/bioinformatics/btu638.

Astorgues-Xerri, L., Riveiro, M.E., Tijeras-Raballand, A., Serova, M., Neuzillet, C., Albert, S., Raymond, E., and Faivre, S. (2014a). Unraveling galectin-1 as a novel therapeutic target for cancer. Cancer Treat Rev 40, 307–319. 10.1016/j.ctrv.2013.07.007.

Astorgues-Xerri, L., Riveiro, M.E., Tijeras-Raballand, A., Serova, M., Rabinovich, G.A., Bieche, I., Vidaud, M., de Gramont, A., Martinet, M., Cvitkovic, E., et al. (2014b). OTX008, a selective small-molecule inhibitor of galectin-1, downregulates cancer cell proliferation, invasion and tumour angiogenesis. Eur J Cancer 50, 2463–2477. 10.1016/j.ejca.2014.06.015.

Bao, S., Wu, Q., McLendon, R.E., Hao, Y., Shi, Q., Hjelmeland, A.B., Dewhirst, M.W., Bigner, D.D., and Rich, J.N. (2006). Glioma stem cells promote radioresistance by preferential activation of the DNA damage response. Nature 444, 756–760. 10.1038/nature05236.

Brooks, M.D., Burness, M.L., and Wicha, M.S. (2015). Therapeutic Implications of Cellular Heterogeneity and Plasticity in Breast Cancer. Cell Stem Cell 17, 260–271. 10.1016/j.stem.2015.08.014.

Camby, I., Le Mercier, M., Lefranc, F., and Kiss, R. (2006). Galectin-1: a small protein with major functions. Glycobiology 16, 137R–157R. o.

Cancer Genome Atlas Research, N. (2008). Comprehensive genomic characterization defines human glioblastoma genes and core pathways. Nature 455, 1061–1068. 10.1038/nature07385.

Carro, M.S., Lim, W.K., Alvarez, M.J., Bollo, R.J., Zhao, X., Snyder, E.Y., Sulman, E.P., Anne, S.L., Doetsch, F., Colman, H., et al. (2010). The transcriptional network for mesenchymal transformation of brain tumours. Nature 463, 318–325. 10.1038/nature08712.

Chen, J., Li, Y., Yu, T.S., McKay, R.M., Burns, D.K., Kernie, S.G., and Parada, L.F. (2012a). A restricted cell population propagates glioblastoma growth after chemotherapy. Nature 488, 522–526. 10.1038/nature11287.

Chen, J., McKay, R.M., and Parada, L.F. (2012b). Malignant glioma: lessons from genomics, mouse models, and stem cells. Cell 149, 36–47. 10.1016/j.cell.2012.03.009.

Chesnelong, C., Hao, X., Cseh, O., Wang, A.Y., Luchman, H.A., and Weiss, S. (2019). SLUG Directs the Precursor State of Human Brain Tumor Stem Cells. Cancers (Basel) 11. 10.3390/cancers11111635.

Chinot, O.L., Wick, W., Mason, W., Henriksson, R., Saran, F., Nishikawa, R., Carpentier, A.F., Hoang-Xuan, K., Kavan, P., Cernea, D., et al. (2014). Bevacizumab plus radiotherapy-temozolomide for newly diagnosed glioblastoma. N Engl J Med 370, 709–722.10.1056/NEJMoa1308345.

Cimino, P.J., Kim, Y., Wu, H.J., Alexander, J., Wirsching, H.G., Szulzewsky, F., Pitter, K., Ozawa, T., Wang, J., Vazquez, J., et al. (2018). Increased HOXA5 expression provides a selective advantage for gain of whole chromosome 7 in IDH wild-type glioblastoma. Genes Dev 32, 512–523. 10.1101/gad.312157.118.

Colaprico, A., Silva, T.C., Olsen, C., Garofano, L., Cava, C., Garolini, D., Sabedot, T.S., Malta, T.M., Pagnotta, S.M., Castiglioni, I., et al. (2016). TCGAbiolinks: an R/Bioconductor package for integrative analysis of TCGA data. Nucleic Acids Res 44, e71. 10.1093/nar/gkv1507.

Cusulin, C., Chesnelong, C., Bose, P., Bilenky, M., Kopciuk, K., Chan, J.A., Cairncross, J.G., Jones, S.J., Marra, M.A., Luchman, H.A., and Weiss, S. (2015). Precursor States of Brain Tumor Initiating Cell Lines Are Predictive of Survival in Xenografts and Associated with Glioblastoma Subtypes. Stem Cell Reports 5, 1–9. 10.1016/j.stemcr.2015.05.010.

de la Iglesia, N., Konopka, G., Puram, S.V., Chan, J.A., Bachoo, R.M., You, M.J., Levy, D.E., Depinho, R.A., and Bonni, A. (2008). Identification of a PTEN-regulated STAT3 brain tumor suppressor pathway. Genes Dev 22, 449–462. 10.1101/gad.1606508.

Dell’albani, P., Rodolico, M., Pellitteri, R., Tricarichi, E., Torrisi, S.A., D’Antoni, S., Zappia, M., Albanese, V., Caltabiano, R., Platania, N., et al. (2014). Differential patterns of NOTCH1-4 receptor expression are markers of glioma cell differentiation. Neuro Oncol 16, 204–216. 10.1093/neuonc/not168.

Ekstrand, A.J., James, C.D., Cavenee, W.K., Seliger, B., Pettersson, R.F., and Collins, V.P. (1991). Genes for epidermal growth factor receptor, transforming growth factor alpha, and epidermal growth factor and their expression in human gliomas in vivo. Cancer Res 51, 2164–2172.

Ekstrand, A.J., Sugawa, N., James, C.D., and Collins, V.P. (1992). Amplified and rearranged epidermal growth factor receptor genes in human glioblastomas reveal deletions of sequences encoding portions of the N- and/or C-terminal tails. Proc Natl Acad Sci U S A 89, 4309–4313. 10.1073/pnas.89.10.4309.

El-Sehemy, A., Selvadurai, H., Ortin-Martinez, A., Pokrajac, N., Mamatjan, Y., Tachibana, N., Rowland, K., Lee, L., Park, N., Aldape, K., et al. (2020). Norrin mediates tumor-promoting and - suppressive effects in glioblastoma via Notch and Wnt. J Clin Invest 130, 3069–3086. 10.1172/JCI128994.

Eyler, C.E., Wu, Q., Yan, K., MacSwords, J.M., Chandler-Militello, D., Misuraca, K.L., Lathia, J.D., Forrester, M.T., Lee, J., Stamler, J.S., et al. (2011). Glioma stem cell proliferation and tumor growth are promoted by nitric oxide synthase-2. Cell 146, 53–66. 10.1016/j.cell.2011.06.006.

Fabregat, A., Jupe, S., Matthews, L., Sidiropoulos, K., Gillespie, M., Garapati, P., Haw, R., Jassal, B., Korninger, F., May, B., et al. (2018). The Reactome Pathway Knowledgebase. Nucleic Acids Res 46, D649–D655. 10.1093/nar/gkx1132.

Fan, Q.W., Cheng, C.K., Gustafson, W.C., Charron, E., Zipper, P., Wong, R.A., Chen, J., Lau, J., Knobbe-Thomsen, C., Weller, M., et al. (2013). EGFR phosphorylates tumor-derived EGFRvIII driving STAT3/5 and progression in glioblastoma. Cancer Cell 24, 438–449. 10.1016/j.ccr.2013.09.004.

Frankish, A., Diekhans, M., Ferreira, A.M., Johnson, R., Jungreis, I., Loveland, J., Mudge, J.M., Sisu, C., Wright, J., Armstrong, J., et al. (2019). GENCODE reference annotation for the human and mouse genomes. Nucleic Acids Res 47, D766–D773. 10.1093/nar/gky955.

Furnari, F.B., Fenton, T., Bachoo, R.M., Mukasa, A., Stommel, J.M., Stegh, A., Hahn, W.C., Ligon, K.L., Louis, D.N., Brennan, C., et al. (2007). Malignant astrocytic glioma: genetics, biology, and paths to treatment. Genes Dev 21, 2683–2710. 10.1101/gad.1596707.

Galli, R., Binda, E., Orfanelli, U., Cipelletti, B., Gritti, A., De Vitis, S., Fiocco, R., Foroni, C., Dimeco, F., and Vescovi, A. (2004). Isolation and characterization of tumorigenic, stem-like neural precursors from human glioblastoma. Cancer Res 64, 7011–7021. 10.1158/0008-5472.CAN-04-1364.

Hara, T., Chanoch-Myers, R., Mathewson, N.D., Myskiw, C., Atta, L., Bussema, L., Eichhorn, S.W., Greenwald, A.C., Kinker, G.S., Rodman, C., et al. (2021). Interactions between cancer cells and immune cells drive transitions to mesenchymal-like states in glioblastoma. Cancer Cell 39, 779–792 e711. 10.1016/j.ccell.2021.05.002.

He, S., Nakada, D., and Morrison, S.J. (2009). Mechanisms of stem cell self-renewal. Annu Rev Cell Dev Biol 25, 377–406. 10.1146/annurev.cellbio.042308.113248.

Heckl, D., Kowalczyk, M.S., Yudovich, D., Belizaire, R., Puram, R.V., McConkey, M.E., Thielke, A., Aster, J.C., Regev, A., and Ebert, B.L. (2014). Generation of mouse models of myeloid malignancy with combinatorial genetic lesions using CRISPR-Cas9 genome editing. Nat Biotechnol 32, 941–946. 10.1038/nbt.2951.

Hsu, Y.L., Wu, C.Y., Hung, J.Y., Lin, Y.S., Huang, M.S., and Kuo, P.L. (2013). Galectin-1 promotes lung cancer tumor metastasis by potentiating integrin alpha6beta4 and Notch1/Jagged2 signaling pathway. Carcinogenesis 34, 1370–1381. 10.1093/carcin/bgt040.

Hu, Y., and Smyth, G.K. (2009). ELDA: extreme limiting dilution analysis for comparing depleted and enriched populations in stem cell and other assays. J Immunol Methods 347, 70–78. 10.1016/j.jim.2009.06.008.

Ignatiadis, N., Klaus, B., Zaugg, J.B., and Huber, W. (2016). Data-driven hypothesis weighting increases detection power in genome-scale multiple testing. Nat Methods 13, 577–580. 10.1038/nmeth.3885.

Jahani-Asl, A., Yin, H., Soleimani, V.D., Haque, T., Luchman, H.A., Chang, N.C., Sincennes, M.C., Puram, S.V., Scott, A.M., Lorimer, I.A., et al. (2016). Control of glioblastoma tumorigenesis by feed-forward cytokine signaling. Nat Neurosci 19, 798–806. 10.1038/nn.4295.

Jensen, K.V., Hao, X., Aman, A., Luchman, H.A., and Weiss, S. (2020). EGFR blockade in GBM brain tumor stem cells synergizes with JAK2/STAT3 pathway inhibition to abrogate compensatory mechanisms in vitro and in vivo. Neurooncol Adv 2, vdaa020. 10.1093/noajnl/vdaa020.

Kim, D., Langmead, B., and Salzberg, S.L. (2015). HISAT: a fast spliced aligner with low memory requirements. Nat Methods 12, 357–360. 10.1038/nmeth.3317.

Kopan, R., and Ilagan, M.X. (2009). The canonical Notch signaling pathway: unfolding the activation mechanism. Cell 137, 216–233. 10.1016/j.cell.2009.03.045.

Korotkevich, G., Sukhov, V., and Sergushichev, A. (2019). Fast gene set enrichment analysis. bioRxiv, 060012. 10.1101/060012.

Kreso, A., and Dick, J.E. (2014). Evolution of the cancer stem cell model. Cell Stem Cell 14, 275–291. 10.1016/j.stem.2014.02.006.

Kros, J.M., Mustafa, D.M., Dekker, L.J., Sillevis Smitt, P.A., Luider, T.M., and Zheng, P.P. (2015). Circulating glioma biomarkers. Neuro Oncol 17, 343–360. 10.1093/neuonc/nou207.

Kwon, A.T., Arenillas, D.J., Worsley Hunt, R., and Wasserman, W.W. (2012). oPOSSUM-3: advanced analysis of regulatory motif over-representation across genes or ChIP-Seq datasets. G3 (Bethesda) 2, 987–1002. 10.1534/g3.112.003202.

Langmead, B., and Salzberg, S.L. (2012). Fast gapped-read alignment with Bowtie 2. Nat Methods 9, 357–359. 10.1038/nmeth.1923.

Lathia, J.D., Mack, S.C., Mulkearns-Hubert, E.E., Valentim, C.L., and Rich, J.N. (2015). Cancer stem cells in glioblastoma. Genes Dev 29, 1203–1217. 10.1101/gad.261982.115.

Lavictoire, S.J., Parolin, D.A., Klimowicz, A.C., Kelly, J.F., and Lorimer, I.A. (2003). Interaction of Hsp90 with the nascent form of the mutant epidermal growth factor receptor EGFRvIII. J Biol Chem 278, 5292–5299. 10.1074/jbc.M209494200.

Le Mercier, M., Mathieu, V., Haibe-Kains, B., Bontempi, G., Mijatovic, T., Decaestecker, C., Kiss, R., and Lefranc, F. (2008). Knocking down galectin 1 in human hs683 glioblastoma cells impairs both angiogenesis and endoplasmic reticulum stress responses. J Neuropathol Exp Neurol 67, 456–469. 10.1097/NEN.0b013e318170f892.

Liberzon, A., Subramanian, A., Pinchback, R., Thorvaldsdottir, H., Tamayo, P., and Mesirov, J.P. (2011). Molecular signatures database (MSigDB) 3.0. Bioinformatics 27, 1739–1740. 10.1093/bioinformatics/btr260.

Liu, F., Hon, G.C., Villa, G.R., Turner, K.M., Ikegami, S., Yang, H., Ye, Z., Li, B., Kuan, S., Lee, A.Y., et al. (2015). EGFR Mutation Promotes Glioblastoma through Epigenome and Transcription Factor Network Remodeling. Mol Cell 60, 307–318. 10.1016/j.molcel.2015.09.002.

Liu, F.T., and Rabinovich, G.A. (2005). Galectins as modulators of tumour progression. Nat Rev Cancer 5, 29–41. 10.1038/nrc1527.

Liu, W., Kim, G.B., Krump, N.A., Zhou, Y., Riley, J.L., and You, J. (2020). Selective reactivation of STING signaling to target Merkel cell carcinoma. Proc Natl Acad Sci U S A 117, 13730–13739. 10.1073/pnas.1919690117.

Lo, H.W., Hsu, S.C., Ali-Seyed, M., Gunduz, M., Xia, W., Wei, Y., Bartholomeusz, G., Shih, J.Y., and Hung, M.C. (2005). Nuclear interaction of EGFR and STAT3 in the activation of the iNOS/NO pathway. Cancer Cell 7, 575–589. 10.1016/j.ccr.2005.05.007.

Love, M.I., Huber, W., and Anders, S. (2014). Moderated estimation of fold change and dispersion for RNA-seq data with DESeq2. Genome Biol 15, 550. 10.1186/s13059-014-0550-8.

Malden, L.T., Novak, U., Kaye, A.H., and Burgess, A.W. (1988). Selective amplification of the cytoplasmic domain of the epidermal growth factor receptor gene in glioblastoma multiforme. Cancer Res 48, 2711–2714.

Navarro, P., Martinez-Bosch, N., Blidner, A.G., and Rabinovich, G.A. (2020). Impact of Galectins in Resistance to Anticancer Therapies. Clin Cancer Res 26, 6086–6101. 10.1158/1078-0432.CCR-18-3870.

Neftel, C., Laffy, J., Filbin, M.G., Hara, T., Shore, M.E., Rahme, G.J., Richman, A.R., Silverbush, D., Shaw, M.L., Hebert, C.M., et al. (2019). An Integrative Model of Cellular States, Plasticity, and Genetics for Glioblastoma. Cell 178, 835–849 e821. 10.1016/j.cell.2019.06.024.

Park, N.I., Guilhamon, P., Desai, K., McAdam, R.F., Langille, E., O’Connor, M., Lan, X., Whetstone, H., Coutinho, F.J., Vanner, R.J., et al. (2017). ASCL1 Reorganizes Chromatin to Direct Neuronal Fate and Suppress Tumorigenicity of Glioblastoma Stem Cells. Cell Stem Cell 21, 209–224 e207. 10.1016/j.stem.2017.06.004.

Phillips, H.S., Kharbanda, S., Chen, R., Forrest, W.F., Soriano, R.H., Wu, T.D., Misra, A., Nigro, J.M., Colman, H., Soroceanu, L., et al. (2006). Molecular subclasses of high-grade glioma predict prognosis, delineate a pattern of disease progression, and resemble stages in neurogenesis. Cancer Cell 9, 157–173. 10.1016/j.ccr.2006.02.019.

Puram, S.V., Yeung, C.M., Jahani-Asl, A., Lin, C., de la Iglesia, N., Konopka, G., Jackson-Grusby, L., and Bonni, A. (2012). STAT3-iNOS Signaling Mediates EGFRvIII-Induced Glial Proliferation and Transformation. J Neurosci 32, 7806–7818. 10.1523/JNEUROSCI.3243-11.2012.

Purow, B.W., Haque, R.M., Noel, M.W., Su, Q., Burdick, M.J., Lee, J., Sundaresan, T., Pastorino, S., Park, J.K., Mikolaenko, I., et al. (2005). Expression of Notch-1 and its ligands, Delta-like-1 and Jagged-1, is critical for glioma cell survival and proliferation. Cancer Res 65, 2353–2363. 10.1158/0008-5472.CAN-04-1890.

Quinlan, A.R. (2014). BEDTools: The Swiss-Army Tool for Genome Feature Analysis. Curr Protoc Bioinformatics 47, 11 12 11–34. 10.1002/0471250953.bi1112s47.

Raizer, J.J., Abrey, L.E., Lassman, A.B., Chang, S.M., Lamborn, K.R., Kuhn, J.G., Yung, W.K., Gilbert, M.R., Aldape, K.A., Wen, P.Y., et al. (2010). A phase II trial of erlotinib in patients with recurrent malignant gliomas and nonprogressive glioblastoma multiforme postradiation therapy. Neuro Oncol 12, 95–103. 10.1093/neuonc/nop015.

Rajakulendran, N., Rowland, K.J., Selvadurai, H.J., Ahmadi, M., Park, N.I., Naumenko, S., Dolma, S., Ward, R.J., So, M., Lee, L., et al. (2019). Wnt and Notch signaling govern self-renewal and differentiation in a subset of human glioblastoma stem cells. Genes Dev 33, 498–510. 10.1101/gad.321968.118.

Raman, V., Martensen, S.A., Reisman, D., Evron, E., Odenwald, W.F., Jaffee, E., Marks, J., and Sukumar, S. (2000). Compromised HOXA5 function can limit p53 expression in human breast tumours. Nature 405, 974–978. 10.1038/35016125.

Ranganathan, P., Weaver, K.L., and Capobianco, A.J. (2011). Notch signalling in solid tumours: a little bit of everything but not all the time. Nat Rev Cancer 11, 338–351. 10.1038/nrc3035.

Reardon, D.A., Nabors, L.B., Mason, W.P., Perry, J.R., Shapiro, W., Kavan, P., Mathieu, D., Phuphanich, S., Cseh, A., Fu, Y., et al. (2015). Phase I/randomized phase II study of afatinib, an irreversible ErbB family blocker, with or without protracted temozolomide in adults with recurrent glioblastoma. Neuro Oncol 17, 430–439. 10.1093/neuonc/nou160.

Rorive, S., Belot, N., Decaestecker, C., Lefranc, F., Gordower, L., Micik, S., Maurage, C.A., Kaltner, H., Ruchoux, M.M., Danguy, A., et al. (2001). Galectin-1 is highly expressed in human gliomas with relevance for modulation of invasion of tumor astrocytes into the brain parenchyma. Glia 33, 241–255. 10.1002/1098-1136(200103)33:3<241::aid-glia1023>3.0.co;2-1.

Schaefer, T.S., Sanders, L.K., and Nathans, D. (1995). Cooperative transcriptional activity of Jun and Stat3 beta, a short form of Stat3. Proc Natl Acad Sci U S A 92, 9097–9101. 10.1073/pnas.92.20.9097.

Schindelin, J., Arganda-Carreras, I., Frise, E., Kaynig, V., Longair, M., Pietzsch, T., Preibisch, S., Rueden, C., Saalfeld, S., Schmid, B., et al. (2012). Fiji: an open-source platform for biological-image analysis. Nat Methods 9, 676–682. 10.1038/nmeth.2019.

Seidel, H.M., Milocco, L.H., Lamb, P., Darnell, J.E., Jr., Stein, R.B., and Rosen, J. (1995). Spacing of palindromic half sites as a determinant of selective STAT (signal transducers and activators of transcription) DNA binding and transcriptional activity. Proc Natl Acad Sci U S A 92, 3041–3045. 10.1073/pnas.92.7.3041.

Sharanek, A., Burban, A., Laaper, M., Heckel, E., Joyal, J.S., Soleimani, V.D., and Jahani-Asl, A. (2020). OSMR controls glioma stem cell respiration and confers resistance of glioblastoma to ionizing radiation. Nat Commun 11, 4116. 10.1038/s41467-020-17885-z.

Singh, S.K., Hawkins, C., Clarke, I.D., Squire, J.A., Bayani, J., Hide, T., Henkelman, R.M., Cusimano, M.D., and Dirks, P.B. (2004). Identification of human brain tumour initiating cells. Nature 432, 396–401. 10.1038/nature03128.

Smedley, D., Haider, S., Durinck, S., Pandini, L., Provero, P., Allen, J., Arnaiz, O., Awedh, M.H., Baldock, R., Barbiera, G., et al. (2015). The BioMart community portal: an innovative alternative to large, centralized data repositories. Nucleic Acids Res 43, W589–598. 10.1093/nar/gkv350.

Soleimani, V.D., Palidwor, G.A., Ramachandran, P., Perkins, T.J., and Rudnicki, M.A. (2013). Chromatin tandem affinity purification sequencing. Nat Protoc 8, 1525–1534. 10.1038/nprot.2013.088.

Somasundaram, K., Reddy, S.P., Vinnakota, K., Britto, R., Subbarayan, M., Nambiar, S., Hebbar, A., Samuel, C., Shetty, M., Sreepathi, H.K., et al. (2005). Upregulation of ASCL1 and inhibition of Notch signaling pathway characterize progressive astrocytoma. Oncogene 24, 7073–7083. 10.1038/sj.onc.1208865.

Stechishin, O.D., Luchman, H.A., Ruan, Y., Blough, M.D., Nguyen, S.A., Kelly, J.J., Cairncross, J.G., and Weiss, S. (2013). On-target JAK2/STAT3 inhibition slows disease progression in orthotopic xenografts of human glioblastoma brain tumor stem cells. Neuro Oncol 15, 198–207. 10.1093/neuonc/nos302.

Stupp, R., Mason, W.P., van den Bent, M.J., Weller, M., Fisher, B., Taphoorn, M.J., Belanger, K., Brandes, A.A., Marosi, C., Bogdahn, U., et al. (2005). Radiotherapy plus concomitant and adjuvant temozolomide for glioblastoma. N Engl J Med 352, 987–996. 10.1056/NEJMoa043330.

Toussaint, L.G., 3rd, Nilson, A.E., Goble, J.M., Ballman, K.V., James, C.D., Lefranc, F., Kiss, R., and Uhm, J.H. (2012). Galectin-1, a gene preferentially expressed at the tumor margin, promotes glioblastoma cell invasion. Mol Cancer 11, 32. 10.1186/1476-4598-11-32.

Venugopal, C., Hallett, R., Vora, P., Manoranjan, B., Mahendram, S., Qazi, M.A., McFarlane, N., Subapanditha, M., Nolte, S.M., Singh, M., et al. (2015). Pyrvinium Targets CD133 in Human Glioblastoma Brain Tumor-Initiating Cells. Clin Cancer Res 21, 5324–5337. 10.1158/1078-0432.CCR-14-3147.

Verhaak, R.G., Hoadley, K.A., Purdom, E., Wang, V., Qi, Y., Wilkerson, M.D., Miller, C.R., Ding, L., Golub, T., Mesirov, J.P., et al. (2010). Integrated genomic analysis identifies clinically relevant subtypes of glioblastoma characterized by abnormalities in PDGFRA, IDH1, EGFR, and NF1. Cancer Cell 17, 98–110. 10.1016/j.ccr.2009.12.020.

Verschuere, T., Toelen, J., Maes, W., Poirier, F., Boon, L., Tousseyn, T., Mathivet, T., Gerhardt, H., Mathieu, V., Kiss, R., et al. (2014). Glioma-derived galectin-1 regulates innate and adaptive antitumor immunity. Int J Cancer 134, 873–884. 10.1002/ijc.28426.

Wheeler, S.E., Suzuki, S., Thomas, S.M., Sen, M., Leeman-Neill, R.J., Chiosea, S.I., Kuan, C.T., Bigner, D.D., Gooding, W.E., Lai, S.Y., and Grandis, J.R. (2010). Epidermal growth factor receptor variant III mediates head and neck cancer cell invasion via STAT3 activation. Oncogene 29, 5135–5145. 10.1038/onc.2009.279.

Wikstrand, C.J., Hale, L.P., Batra, S.K., Hill, M.L., Humphrey, P.A., Kurpad, S.N., McLendon, R.E., Moscatello, D., Pegram, C.N., Reist, C.J., and, et al. (1995). Monoclonal antibodies against EGFRvIII are tumor specific and react with breast and lung carcinomas and malignant gliomas. Cancer Res 55, 3140–3148.

Yan, J., Enge, M., Whitington, T., Dave, K., Liu, J., Sur, I., Schmierer, B., Jolma, A., Kivioja, T., Taipale, M., and Taipale, J. (2013). Transcription factor binding in human cells occurs in dense clusters formed around cohesin anchor sites. Cell 154, 801–813. 10.1016/j.cell.2013.07.034.

Zhang, C., Hai, L., Zhu, M., Yu, S., Li, T., Lin, Y., Liu, B., Zhou, X., Chen, L., Zhao, P., et al. (2017). Actin cytoskeleton regulator Arp2/3 complex is required for DLL1 activating Notch1 signaling to maintain the stem cell phenotype of glioma initiating cells. Oncotarget 8, 33353–33364. 10.18632/oncotarget.16495.

Zhang, Y., Liu, T., Meyer, C.A., Eeckhoute, J., Johnson, D.S., Bernstein, B.E., Nusbaum, C., Myers, R.M., Brown, M., Li, W., and Liu, X.S. (2008). Model-based analysis of ChIP-Seq (MACS). Genome Biol 9, R137. 10.1186/gb-2008-9-9-r137.

