## Supplementary Figures for "Transcriptional Control of Brain Tumour Stem Cells by a Carbohydrate Binding Protein"

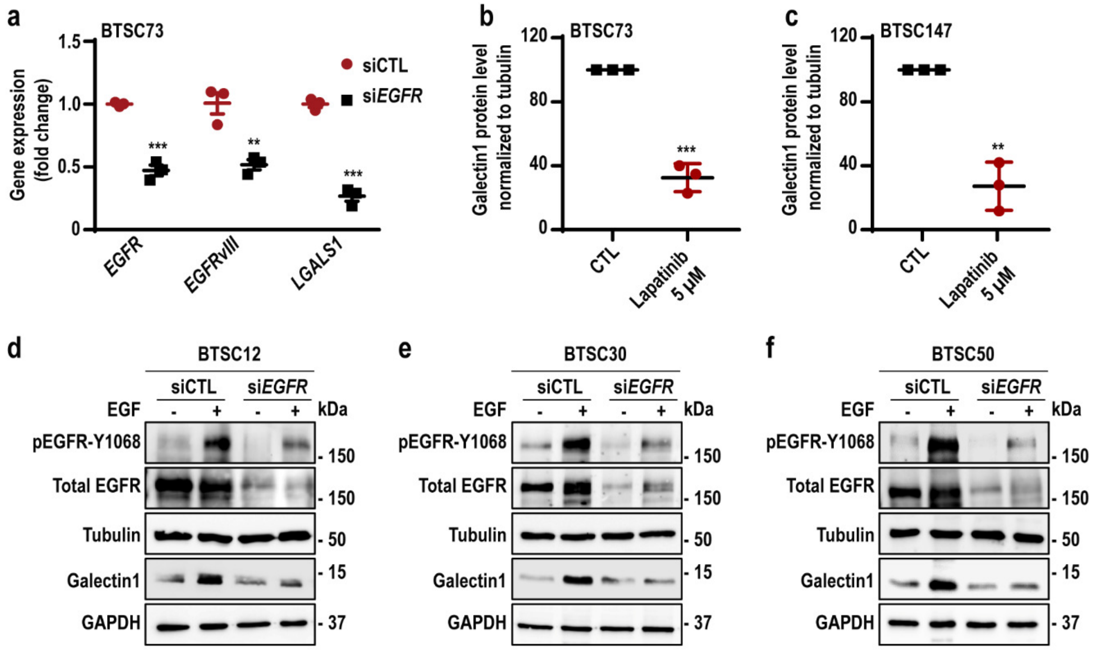


**Figure S1. Galectin1 is upregulated in an EGFRvIII-dependent manner in patient-derived BTSCs.** Related to Figure 1.

(**a**) BTSC73 was electroporated with si*EGFR*/*EGFRvIII* or siCTL. *LGALS1*, *EGFRvIII* and *EGFR* mRNA expression were evaluated by RT-qPCR. GUSB was used as a housekeeping gene. (**b-c**) Densitometric quantification of galectin1 protein expression level normalized to tubulin in lapatinib-treated BTSC73 (b) and BTSC147 (c) are shown (accompanying representative blots in Figure 1e-f). (**d-f**) BTSC12 (d), BTSC30 (e), and BTSC50 (f) that lack EGFRvIII mutation, were electroporated with si*EGFR* or siCTL. Cells were treated with 100 ng/ml of EGF and subjected to immunoblotting using antibodies indicated on the blots. Data are presented as the mean ± SEM, n = 3. Unpaired two-tailed *t*-test (a, b and c), $^{**}p < 0.01$, $^{***}p < 0.001$.

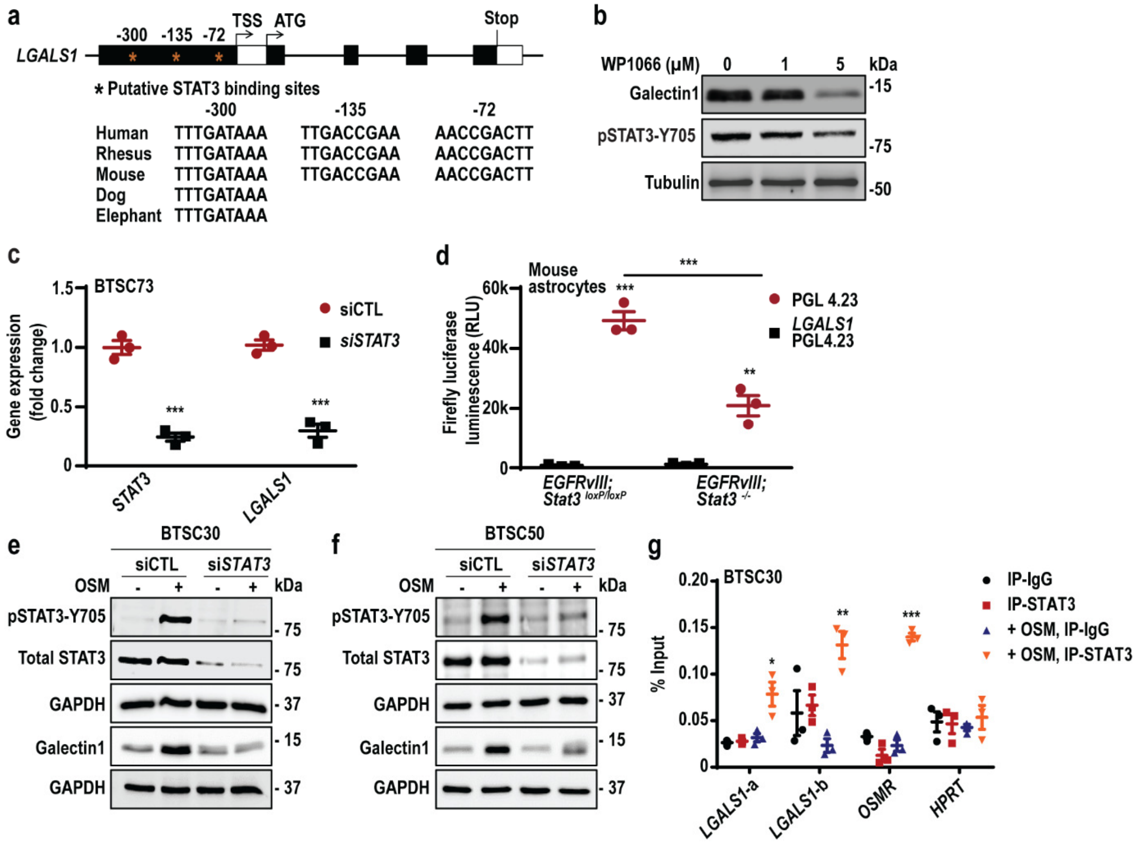


**Figure S2. *LGALS1* is a direct transcriptional target of STAT3.** Related to Figure 1. (**a**) *LGALS1* promoter harbours putative STAT3 binding motifs at positions -300, -135 and -72 bp upstream of the *LGALS1* transcriptional start site (TSS), conserved across different species. (**b**) BTSC73 was treated with 1 or 5 µM of the STAT3 inhibitor, WP1066, or vehicle control (DMSO). Cell lysates were subjected to immunoblotting using antibodies to galectin1 and pSTAT3-Y705. Tubulin was used as a loading control. (**c**) BTSC73 was electroporated with siCTL or si*STAT3*. mRNA levels of *STAT3* and *LGALS1* were evaluated by RT-qPCR. GUSB was used as a housekeeping gene. (**d**) Luciferase reporter assay was performed in *EGFRvIII*;*Stat3*$^{loxP/loxP}$ and *EGFRvIII*;*Stat3*$^{-/-}$ mouse astrocytes. (**e-f**) BTSCs that lack the EGFRvIII mutation were electroporated with si*STAT3* or siCTL, treated with 10 ng/ml OSM for 3h and subjected to immunoblotting analysis using antibodies indicated on the blots. (**g**) BTSC30 was treated with 10 ng/ml OSM then subjected to ChIP using an antibody to STAT3 or IgG control followed by qPCR analysis using two different pairs of primers, *LGALS1*-a and *LGALS1*-b. *OSMR*, and *HPRT* loci were used as positive and negative controls, respectively. Data are presented as the mean ± SEM, n = 3. Unpaired two-tailed *t*-test (c and g); one-way ANOVA followed by Tukey's test (d), *p < 0.05, **p < 0.01, ***p < 0.001.

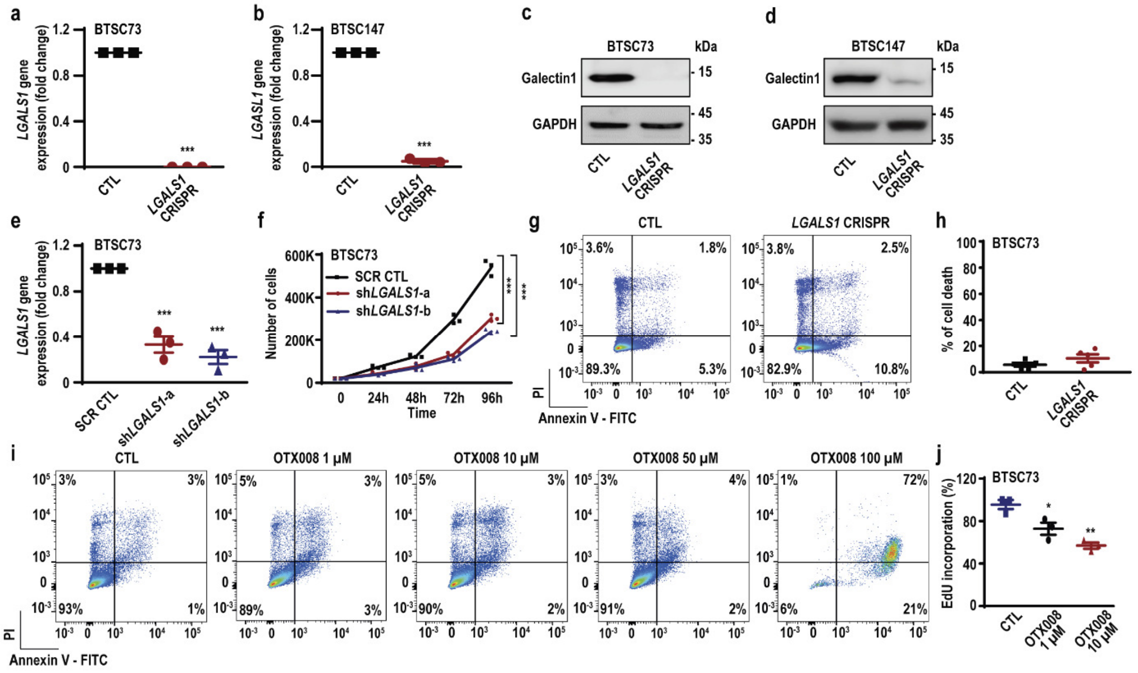


**Figure S3. Galectin1 inhibition suppresses BTSC growth without induction of cell death.** Related to Figure 2.

(**a-d**) *LGALS1* KO was confirmed by subjecting *LGALS1* CRISPR and CTL BTSCs to RT-qPCR (a-b) and immunoblotting analyses (c-d). (**e**) *LGALS1* KD was confirmed by subjecting BTSC73 transduced with two different *LGALS1* shRNA (sh*LGALS1*-a and sh*LGALS1*-b) and scramble shRNA control (SCR CTL) to RT-qPCR analysis. GUSB was used as a housekeeping gene. (**f**) Population growth curves for *LGALS1* KD and control BTSC73 are shown. (**g-h**) Apoptosis analysis was performed in *LGALS1* CRISPR and CTL BTSC73 by flow cytometry following annexin V and PI double staining. Representative scatter plots of flow cytometry analyses are shown (g). The percentage of cell death (annexin V positive cells) is presented in the histogram (h). (**i**) Apoptosis analysis was performed by flow cytometry following annexin V and PI double staining in BTSC73 treated with different concentrations of OTX008. Representative scatter plots of flow cytometry analyses are shown. (**j**) EdU cell proliferation assay was performed 22h following treatment of BTSC73 with 1 or 10 µM of OTX008. Data are presented as the mean ± SEM, $n \geq 3$. Unpaired two-tailed *t*-test (a and b); one-way ANOVA followed by Dunnett's test (e, f and j), $*p < 0.05$, $**p < 0.01$, $***p < 0.001$.

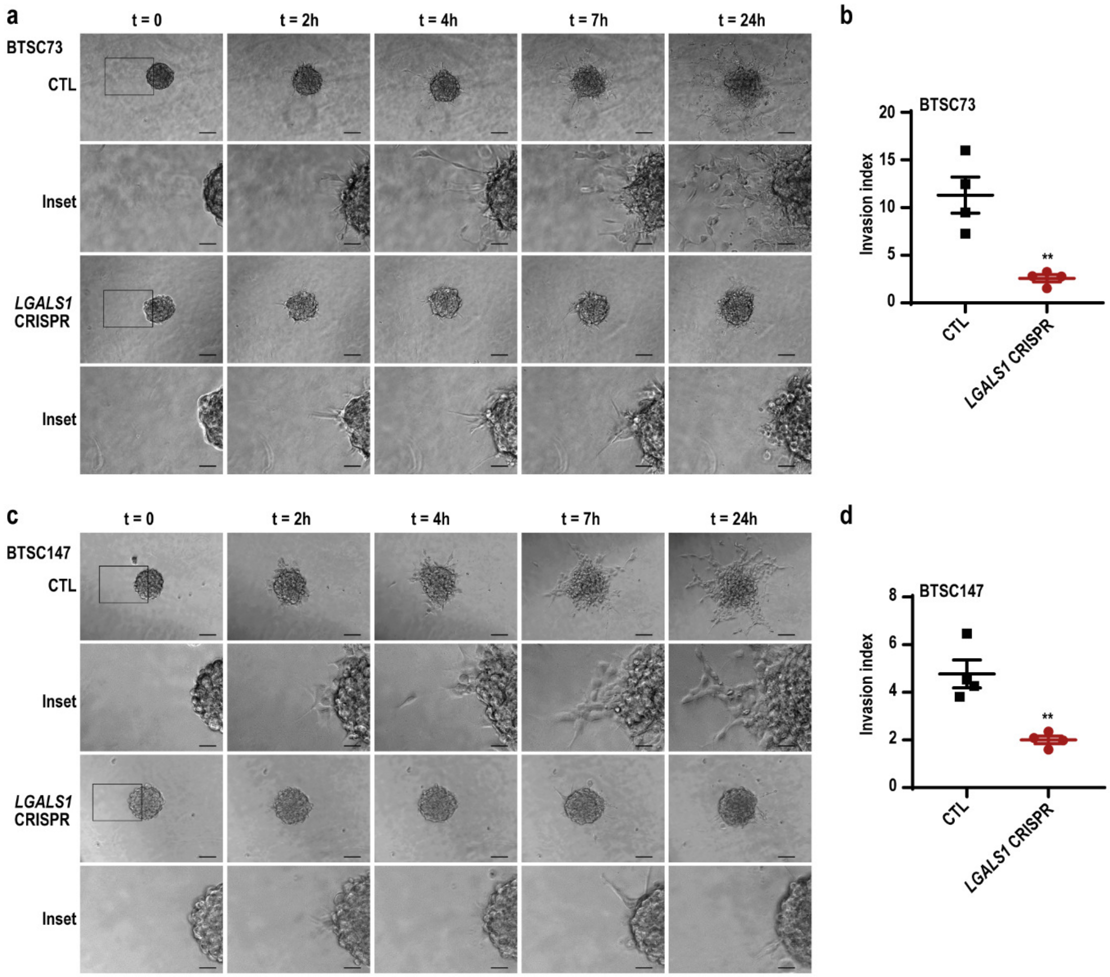


**Figure S4. Galectin1 promotes invasion of EGFRvIII-expressing BTSCs.** Related to Figure 2.

(**a-d**) *LGALS1* CRISPR or CTL BTSC73 and BTSC147 were subjected to collagen type I invasion assay. Representative phase-contrast time-lapse imaging of BTSC73 (a) and BTSC147 (c) spheres in collagen I gels are shown. Scale bar = 100 µm. Inset scale bar = 35 µm. Invasion index was calculated for BTSC73 (b) and BTSC147 (d) after 24 h of culturing BTSC spheres in collagen I gels. Data are presented as the mean ± SEM, n = 4. Unpaired two-tailed *t*-test (b and d), **p < 0.01.

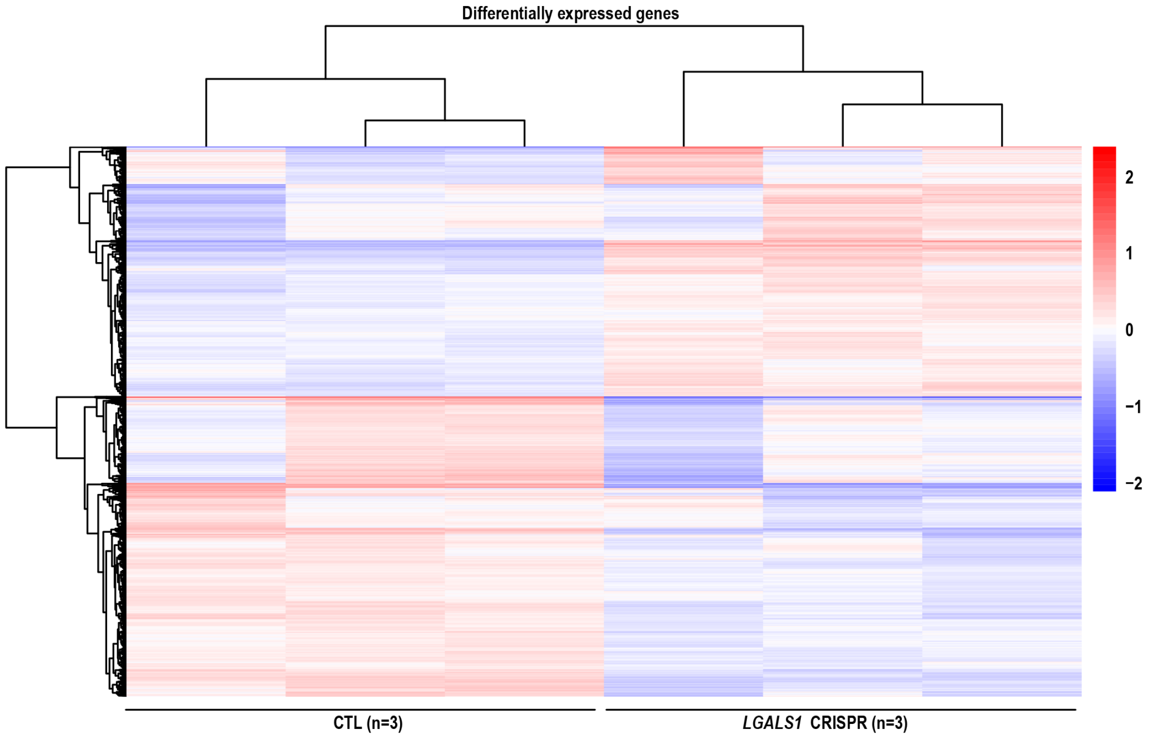


**Figure S5. Gene expression profiling of *LGALS1* CRISPR BTSCs.** Related to Figure 4. *LGALS1* CRISPR and CTL BTSCs were subjected to RNA-sequencing analysis. Heatmap illustration of hierarchical clustering of the overall gene expression profiles is shown. Colour gradient: red, high expression; blue, low expression**.**

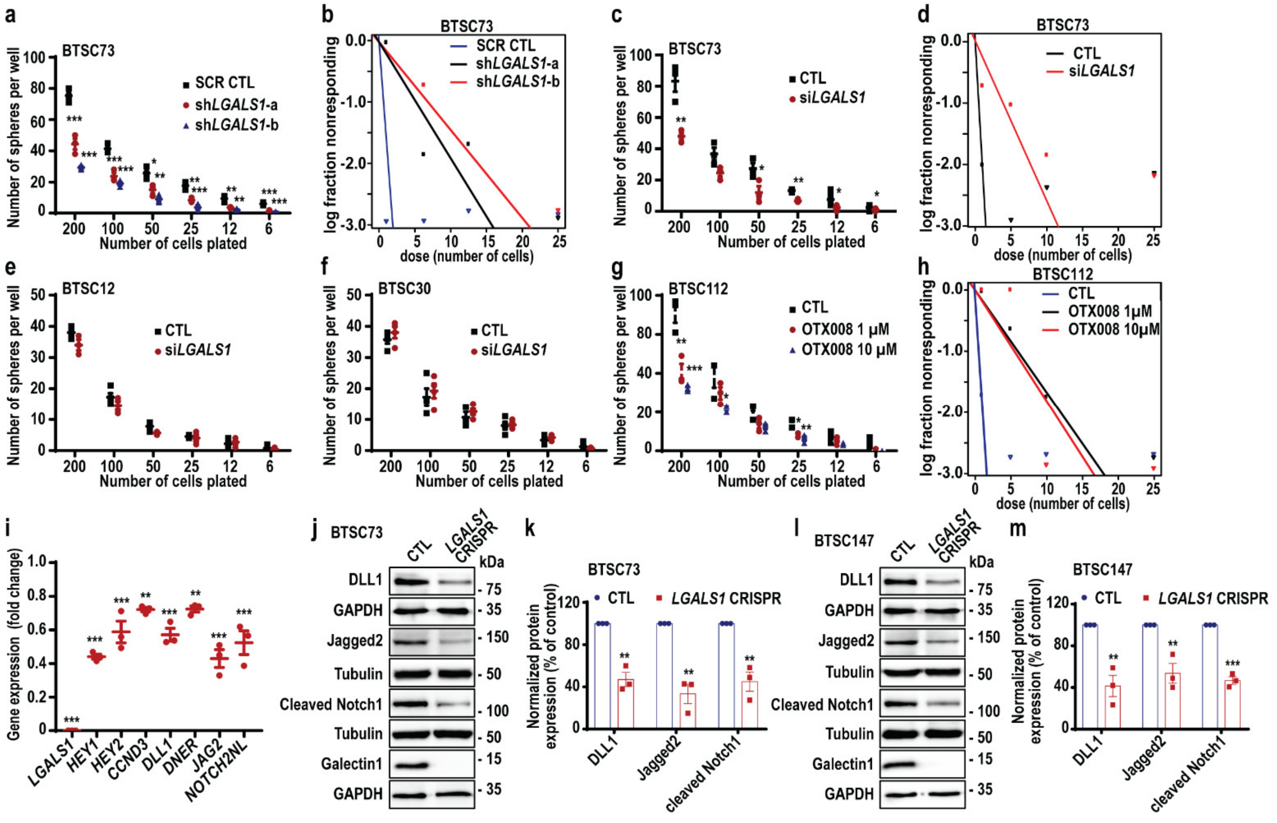


**Figure S6. Galectin1 controls self-renewal of BTSCs.** Related to Figure 5.

(**a-b**) EGFRvIII-expressing BTSC73 was transduced with sh*LGALS1*-a, sh*LGALS1*-b or SCR CTL, and subjected for LDA and ELDA analysis. (**c-d**) EGFRvIII-expressing BTSC73 was electroporated with si*LGALS1* or siCTL and subjected to LDA and ELDA analysis. (**e-f**) BTSC12 (e) and BTSC30 (f) that do not harbour EGFRvIII mutation were electroporated with si*LGALS1* or siCTL and subjected to LDA analysis. (**g-h**) EGFRvIII-expressing BTSC112 was subjected to LDA and ELDA analysis following treatment with 1 or 10 μM OTX008. (**i**) *LGALS1* CRISPR and CTL BTSC73 were subjected to RT-qPCR analysis to assess the expression of Notch signalling related genes. GUSB was used as a housekeeping gene. (**j-m**) EGFRvIII-expressing *LGALS1* CRISPR and CTL BTSC73 (j-k) and BTSC147 (l-m) were analyzed by immunoblotting using antibodies indicated on the blots. Tubulin and GAPDH were used as loading controls. Densitometric quantification of protein expression levels of DLL1 (normalized to GAPDH), Jagged2 (normalized to tubulin) and cleaved Notch1 (normalized to tubulin) in BTSC73 (k) and BTSC147 (m) are shown. Data are presented as the mean ± SEM, n = 3. One-way ANOVA followed by Dunnett's test (a, g and i); unpaired two-tailed *t*-test (c, k and m), *p < 0.05, **p < 0.01, ***p < 0.001.

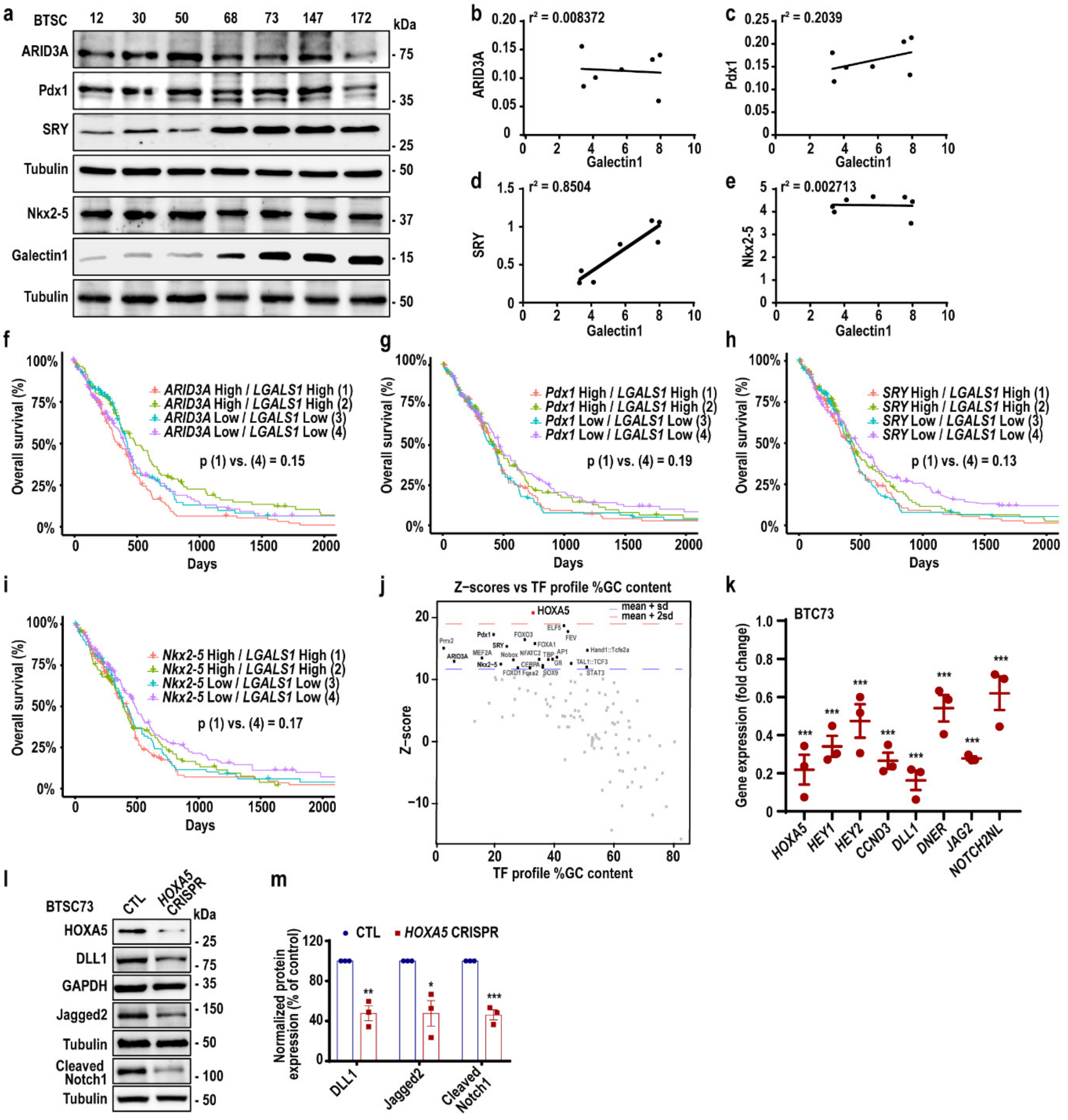

**Figure S7. Screening for transcription factors that may cooperate with galectin1.** Related to Figure 7.

(**a**) Patient-derived BTSCs that harbour EGFRvIII mutation or lack the mutation were analyzed by immunoblotting using antibodies indicated on the blots. Tubulin was used as a loading control. (**b-e**) Pearson analyses were conducted on the densitometric values of protein expression for ARID3A (b), Pdx1 (c), SRY (d) and Nkx2-5 (e) and their correlation with galectin1 expression. (**f-i**) KM survival plots were generated via clustering off glioblastoma patients based on the expression level of *LGALS1* and each of the ARID3A (f), Pdx1 (g), SRY (h) and Nkx2-5 (i). (**j**) *LGALS1*-downregulated genes from RNA-seq analysis were subjected to enrichment analysis of TF binding motifs using oPOSSUM-3 software. High-scoring or over-represented TF binding site profiles were computed as having z-scores above the mean + 2 x standard deviation (red dotted line). (**k**) *HOXA5* CRISPR and CTL BTSC73 were subjected to RT-qPCR to assess the expression levels of Notch signalling related genes. GUSB was used as a housekeeping gene. (**l**) EGFRvIII-expressing *HOXA5* CRISPR and CTL BTSC73 were analyzed by immunoblotting using antibodies indicated on the blots. Tubulin and GAPDH were used as loading controls. (**m**) Densitometric quantification of protein expression levels of DLL1 (normalized to GAPDH), Jagged2 (normalized to tubulin) and cleaved Notch1 (normalized to tubulin). Data are presented as the mean ± SEM, n = 3. Log-rank test (f, g, h and i); one-way ANOVA followed by Dunnett's test (k); unpaired two-tailed *t*-test (m), $*p < 0.05$, $**p < 0.01$, $***p < 0.001$.

**Table S1: Oligonucleotide information, related to STAR Methods**

| | |
|---|---|
| **gRNAs primers used for CRISPR/Cas9 system** | |
| gRNA-1-*LGALS1* | caccGGAGAGTGCCTTCGAGTGCG |
| gRNA-1-*LGALS1*-RC | aaacCGCACTCGAAGGCACTCTCC |
| gRNA-2-*LGALS1* | caccGCCTCCAGGTTGAGGCGGTT |
| gRNA-2-*LGALS1*-RC | aaacAACCGCCTCAACCTGGAGGC |
| gRNA-1-*HOXA5* | caccGTAGCCGTAGCCGTACCTGC |
| gRNA-1-*HOXA5*-RC | aaacGCAGGTACGGCTACGGCTAC |
| gRNA-2-*HOXA5* | caccGCACGCAGGGACACACCGCT |
| gRNA-2-*HOXA5*-RC | aaacAGCGGTGTGTCCCTGCGTGC |
| **Screening primers used for CRISPR/Cas9 system** | |
| *LGALS1* External-Fwd | CTTGGCTTGGTCAGAGGATGC |
| *LGALS1* External-Rev | TTCAGAGGGAGCAGAGGCAG |
| *LGALS1* Internal-Fwd | TCAAGAATCAAGCGAGCCC |
| *LGALS1* Internal-Rev | CAGTATCCCATGAACGCACC |
| *HOXA5* External-Fwd | AGGGTGCTATAGACGCACAAA |
| *HOXA5* External-Rev | AAAACAGCCAGACTTGGACAAT |
| *HOXA5* Internal-Fwd | ACCCACATCAGCAGCAGAG |
| *HOXA5* Internal-Rev | GGTCGCGTGGATTTAGAAAA |
| **qPCR primers** | |
| *GUSB*-Fwd | GCGTTCCTTTTGCGAGGAGA |
| *GUSB*-Rev | GGTGGTATCAGTCTTGCTCAA |
| *LGALS1*-Fwd | CACTTCAACCCTCGCTTCA |
| *LGALS1*-Rev | CGTCAGCTGCCATGTAGTT |
| *HOXA5*-Fwd | TCTCGTTGCCCTAATTCATCTTTT |
| *HOXA5*-Rev | CATTCAGGACAAAGAGATGAACAGA |
| *PIM2*-Fwd | GTGGCTGTGCCAAACTCATT |
| *PIM2*-Rev | ATGCCCAGTGACCAGACAGT |
| *E2F7*-Fwd | GAAAGCACCAAAGAGCCTTCT |
| *E2F7*-Rev | AAGACCATGCAAGGGACACT |
| *NEDD9*-Fwd | TGACTGTAGCAGCAGTGATGG |
| *NEDD9*-Rev | TGTTCCAGCTGCATCTTGTT |
| *MCM5*-Fwd | CAGAGGCAGATGTGGAGGAG |
| *MCM5*-Rev | GCTTGAGCTGCTTCTCGATG |
| *KLHL22*-Fwd | CCACAATGACCTGAATGCTG |

| | |
|---|---|
| *KLHL22*-Rev | TCAGGTAATCCTCCCCTCTG |
| *NDC80*-Fwd | CTGTTAACCAGGGGCTCAGT |
| *NDC80*-Rev | GACCCAACATGTGTAGCAACC |
| *GSPT2*-Fwd | CAAAGATATGGGCACTGTGG |
| *GSPT2*-Rev | GTTTTCACCTGGGGCTACAA |
| *HIRA*-Fwd | ACGCACGGTACCTCGTAAAC |
| *HIRA*-Rev | TGTTGACTCCCACTGGCTTC |
| *SMARCA4*-Fwd | GATGACAGTGAAGGCGAGGA |
| *SMARCA4*-Rev | GGCCAAGCTTGATCTTCACTT |
| *CENPM*-Fwd | TCTTGGGGAAGGTGTGTTTC |
| *CENPM*-Rev | TAGAGCAGGGGGCTTTGATA |
| *GTSE1*-Fwd | CAGAAGTAGCTCGGGAGGAA |
| *GTSE1*-Rev | CTTGCAGCATCTGGAGTGAC |
| *MAD2L2*-Fwd | GCTGTACCTTCACAGTCCTGGT |
| *MAD2L2*-Rev | ATGTCCGACGTCATGGTTTT |
| *NEK6*-Fwd | GACGCCCTACTACATGTCACC |
| *NEK6*-Rev | TGGCACAGGGAGAAGAGATT |
| *SKP2*-Fwd | ACCTTTCTGGGTGTTCTGGA |
| *SKP2*-Rev | CTGGGTGATGGTCTCTGACA |
| *CCND3*-Fwd | ATTTCCTGGCCTTCATTCTG |
| *CCND3*-Rev | CGGGTACATGGCAAAGGTAT |
| *CDK6*-Fwd | CATTCAAAATCTGCCCAACC |
| *CDK6*-Rev | GGTGGGAATCCAGGTTTTCT |
| *CDKN2D*-Fwd | GTCATGATGTTTGGCAGCAC |
| *CDKN2D*-Rev | CGTCATGGACTGGACTGGTA |
| *DNER*-Fwd | TAATGAGTGCCTCTCCGCTC |
| *DNER*-Rev | TAAACCCGGGTGCACAGATG |
| *DLL1*-Fwd | AGCTACACTTGCTCTTGCCG |
| *DLL1*-Rev | GACCCCCGTTAAAGCAAGGG |
| *NOTCH2NLA*-Fwd | GTGTCTCGACCTTGCCTGAA |
| *NOTCH2NLA*-Rev | CTCACACTTCTGCCCTGTGA |
| *HEY2*-Fwd | GGCGTCGGGATCGGATAAAT |
| *HEY2*-Rev | TTACCCCCTGTTGCCTGAAG |
| *HEY1*-Fwd | TAATTGAGAAGCGCCGACGA |
| *HEY1*-Rev | GGTCATCTGCAGGATCTCGG |

| | |
|---|---|
| *JAG2*-Fwd | GAGCTGGAACGAGACGAGTG |
| *JAG2*-Rev | CCAAAGTCATCAGGGCAGG |
| **ChIP-PCR primers** | |
| *OSMR*-Fwd | GACTGAAGGGAGGGAATTCCTGT |
| *OSMR*-Rev | CAATTTCCCGTCTTGCTG |
| *LGALS1*-a-Fwd | ACTTGTGGGCCTAGCTCATC |
| *LGALS1*-a-Rev | TTCTCCATCCCTCTCCA |
| *LGALS1*-b-Fwd | GCCACTCTGATTGGTCACCT |
| *LGALS1*-b-Rev | ACGCTCCCACCCTTTTAACT |
| *HPRT*-Fwd | CGGTAGGTTTGGGAATCA |
| *HPRT*-Rev | CAGTTTGCAGGCTCACTA |
| *ARIDA5B*-Fwd | GCGCTGGGTTATATAAACACATTA |
| *ARIDA5B*-Rev | AAATGCGAGAAGCGAGTCTG |
| *DSCAM*-Fwd | AAGGGGCTATATGTTTGGGATT |
| *DSCAM*-Rev | CTCCTTCCAAATCCTTGCTG |
| *EDN1*-Fwd | GGGCAGGTTTAGCAAAGGTC |
| *EDN1*-Rev | TTAGTCACCAACAGGCAACG |
| *KCNK2*-Fwd | CGTGGATGCTTCGTGTGTAA |
| *KCNK2*-Rev | TTCAGGAAGAAATTCCCTGATT |
| *ACSS3*-Fwd | TTGAATATATCTCCTCTTATGACCAC |
| *ACSS3*-Rev | AGATCAGCTTTTTGCTTCTTTG |
| *HBB*-Fwd | CTGTTTGAGGTTGCTAGTG |
| *HBB*-Rev | TCATCACTTAGACCTCACC |
| **Dual-luciferase reporter assay primers** | |
| *LGALS1*-Fwd | CTCAGCCATCTTCTCTGGGC |
| *LGALS1*-Rev | AGTTAAAAGGGTGGGAGCGT |
